# N-Myc and STAT Interactor is an endometriosis suppressor

**DOI:** 10.1101/2024.05.08.593227

**Authors:** Yuri Park, Xiaoming Guan, Sang Jun Han

## Abstract

In patients with endometriosis, refluxed endometrial fragments evade host immunosurveillance, developing into endometriotic lesions. However, the mechanisms underlying this evasion have not been fully elucidated. N-Myc and STAT Interactor (NMI) have been identified as key players in host immunosurveillance, including interferon (IFN)-induced cell death signaling pathways. NMI levels are markedly reduced in the stromal cells of human endometriotic lesions due to modulation by the Estrogen Receptor beta/Histone Deacetylase 8 axis. Knocking down NMI in immortalized human endometrial stromal cells (IHESCs) led to elevated RNA levels of genes involved in cell-to-cell adhesion and extracellular matrix signaling following IFNA treatment. Furthermore, NMI knockdown inhibited IFN-regulated canonical signaling pathways, such as apoptosis mediated by Interferon Stimulated Gene Factor 3, and necroptosis upon IFNA treatment. In contrast, NMI knockdown with IFNA treatment activated non-canonical IFN-regulated signaling pathways that promote proliferation, including β-Catenin and AKT signaling. Moreover, NMI knockdown in IHESCs stimulated ectopic lesions’ growth in mouse endometriosis models. Therefore, NMI is a novel endometriosis suppressor, enhancing apoptosis and inhibiting proliferation and cell adhesion of endometrial cells upon IFN exposure.

## Introduction

Endometriosis is a medical condition in which endometrial tissues grow outside the uterus, such as on the ovaries and in the peritoneal cavity (Zondervan, Becker et al., 2020). Approximately 10% of reproductive-aged women suffer from endometriosis. Symptoms include pelvic or abdominal pain, excessive menstrual flow, pain during urination or bowel movements, and infertility (Zondervan et al., 2020). Conservative surgery is the first-line treatment for endometriosis; however, it cannot prevent a relapse of the condition (Saunders & Horne, 2021). Due to the estrogen dependence of endometriosis, anti-estrogen drugs have been used to alleviate symptoms, suppress post-operative recurrence, and enhance the quality of life for those affected (Lyons, Chew et al., 2006, Zakhari, Delpero et al., 2021). However, hormonal therapy can lead to adverse effects, including post-menopausal symptoms and unintended consequences in other estrogen-responsive organs, such as the bones and brain (Saunders & Horne, 2021, Tosti, Biscione et al., 2017). Thus, new therapeutic approaches for endometriosis based on the discovery of a new molecular etiology are highly needed.

What causes endometriosis? Several hypotheses attempt to answer this question (Sourial, Tempest et al., 2014). Among these, the retrograde menstruation hypothesis is a widely accepted explanation for the progression of endometriosis (Sampson, 1925, Zondervan et al., 2020).

However, while 90% of reproductive-aged women experience retrograde menstruation, only 10% of them are diagnosed with endometriosis (Halme, Hammond et al., 1984). Therefore, other risk factors are also involved in initiating and progressing endometriosis along with retrograde menstruation. Previous studies have proposed that aberrant Estrogen Receptor beta (ERβ) expression in endometrial tissue might be a key driver in the development of endometriosis among reproductive-aged women who have experienced retrograde menstruation (Bulun, 2009, Bulun, Cheng et al., 2010, Monnin, Fattet et al., 2023). Hypomethylation of the ERβ promoter leads to the upregulation of ERβ in endometriotic lesions (Xue, Lin et al., 2007). ERβ promotes the proliferation of endometriotic stromal cells by transcriptionally upregulating the Ras-like and estrogen-regulated growth inhibitor (RERG) in conjunction with prostaglandin (Monsivais, Dyson et al., 2014). Additionally, ERβ interacts with TNFα-induced apoptosis complexes and inflammasomes, promoting endometriotic cell proliferation, adhesion, and invasion (Han, Jung et al., 2015a). Interestingly, ERβ significantly suppresses the Interferon (IFN)-mediated cell death pathways, promoting the growth of endometriotic lesions in mice with endometriosis (Han, Lee et al., 2019a). The aberrant expression of ERβ, in conjunction with retrograde menstruation, might contribute to the onset of endometriosis. However, the key question of how ERβ downregulates Interferon (IFN)-mediated cell death pathways is not addressed.

During retrograde menstruation, refluxed endometrial fragments activate host immune surveillance systems and elevate proinflammatory cytokines (such as TNFα and IFNs) in the pelvic area (Park & Han, 2022). IFNA induces cell death in malignant cells through various mechanisms (Shi, Yao et al., 2022). For example, IFNα activates the caspase-3-mediated intrinsic apoptotic pathway as well as the caspase-4-mediated endoplasmic reticulum (ER) stress-induced apoptosis in HeLa cervical cancer cells (Shi, Cao et al., 2016). In hepatocellular carcinoma cells, IFNA treatment enhances the expression of promyelocytic leukemia (PML) and tumor necrosis factor-related apoptosis-inducing ligand (TRAIL), activating the extrinsic apoptotic pathways that lead to cell death (Herzer, Hofmann et al., 2009). In the canonical IFNA pathway, Janus Kinase 1 (JAK1) and Tyrosine Kinase 2 (TYK2) phosphorylate Signal Transducers and Activators of Transcription 1 (STAT1) and 2 (STAT2). These phosphorylated forms subsequently dimerize and, in collaboration with Interferon Regulatory Factor 9 (IRF9), bind to the Interferon-Stimulated Response Element (ISRE) (Mazewski, Perez et al., 2020). In endometriosis, type I IFN (including IFNA and IFNB) is dysregulated (Park & Han, 2022, Pia, Anna et al., 2012). The mRNA levels of IFNα and its receptor 2 (IFNAR2) are elevated in the eutopic endometrium of endometriosis patients compared to healthy women (Kao, Germeyer et al., 2003). A key mediator of the type I IFN pathway, JAK1, is significantly upregulated in endometriotic lesions compared to the eutopic endometrium of endometriosis patients. This suggests that the IFNA/IFNAR2/JAK1 signaling pathway is essential in endometriosis progression. (Matsuzaki, Canis et al., 2006, Park & Han, 2022). However, the transition from IFNA-mediated cell death signaling to IFNA-mediated growth stimulation signaling in endometriotic lesions, as opposed to the normal endometrium, remains unclear in the context of endometriosis progression.

Our microarray analysis demonstrated a significant reduction in N-Myc and STAT interactor (NMI) levels within endometriotic lesions, compared to the normal endometrium of mice with endometriosis (Han, Lee et al., 2019b). NMI is an interferon-inducible gene, with its promoter region containing an ISRE (Lebrun, Shpall et al., 1998, Xu, Chai et al., 2018). NMI interacts with all STATs except STAT2 (Zhu, John et al., 1999). Although NMI does not possess an intrinsic transcriptional activation domain, it enhances IL-2 and IFNG signaling. This enhancement occurs through the facilitation of STAT5 and STAT1 association with the transcriptional coactivators CBP/p300 (Zhu et al., 1999). NMI exhibits anti-tumor activity, as evidenced by its significantly low expression levels in breast cancer cells. Furthermore, the downregulation of NMI is associated with the promotion of human lung adenocarcinoma, and low expression levels of NMI correlate with a poor prognosis in lung cancer (Fillmore, Mitra et al., 2009, Wang, Zou et al., 2017). In glioblastoma, however, NMI exhibits oncogenic activity, as its expression is positively correlated with poor prognosis. This is because NMI promotes cell cycle progression and proliferation through STAT1 (Meng, Chen et al., 2015).

Although the precise role of NMI in the progression of endometriosis remains unclear, our study has identified NMI’s function within this context: NMI acts as a suppressor of endometriosis.

The downregulation of NMI, mediated by the Estrogen Receptor beta/Histone Deacetylase 8 axis, triggers key endometriotic phenotypes—such as anti-apoptosis, hyperproliferation, and increased cell adhesion—in endometrial cells, leading to the development of endometriotic lesions.

## Results

### NMI levels are reduced in stromal cells of human endometriotic lesions

We utilized primary human endometrial stromal cells derived from ectopic lesions in patients with endometriosis and from normal endometrium that we have isolated (Mohankumar, Li et al., 2020). We subsequently measured NMI levels in both types of endometrial cells using Western blot analysis. NMI levels were found to be lower in human endometrial stromal cells from endometriosis patients compared to those in normal endometrial stromal cells (Fig. 1A, B). To confirm that these endometrial stromal cells originated from endometriosis patients, we assessed ERβ levels in the cells, given that ERβ levels are known to be elevated in endometriotic lesions compared to normal endometrium (Bulun et al., 2010, Han, Jung et al., 2015b). In line with these observations, ERβ levels were significantly higher in stromal cells from endometriosis patients compared to normal endometrial stromal cells (Fig. 1A, C). Furthermore, NMI expression was found to be inversely correlated with ERβ levels when comparing human endometriotic stromal cells with normal endometrial stromal cells. Next, we evaluated NMI levels in ectopic lesions from human endometriosis patients compared to normal endometrium using the GSE25628 dataset from the Gene Expression Omnibus (GEO) database, which is specific to endometriosis patients (Fig. 1D). We found that NMI mRNA levels in human ectopic lesions were significantly lower than those in normal endometrium. Furthermore, we examined NMI protein levels in human ectopic lesions relative to normal endometrium using immunohistochemistry (IHC). The H-Score for NMI indicated that NMI protein levels in the epithelium of ectopic lesions were similar to those in the normal endometrial epithelium (Fig. 1E, F). However, NMI protein levels in the stromal cells of ectopic lesions were significantly lower than those in normal endometrium (Fig. 1G). This significant reduction in stromal NMI levels within endometriotic lesions, compared to normal endometrium, suggests that the decline in stromal NMI may be associated with the progression of endometriosis.

**Fig 1.**
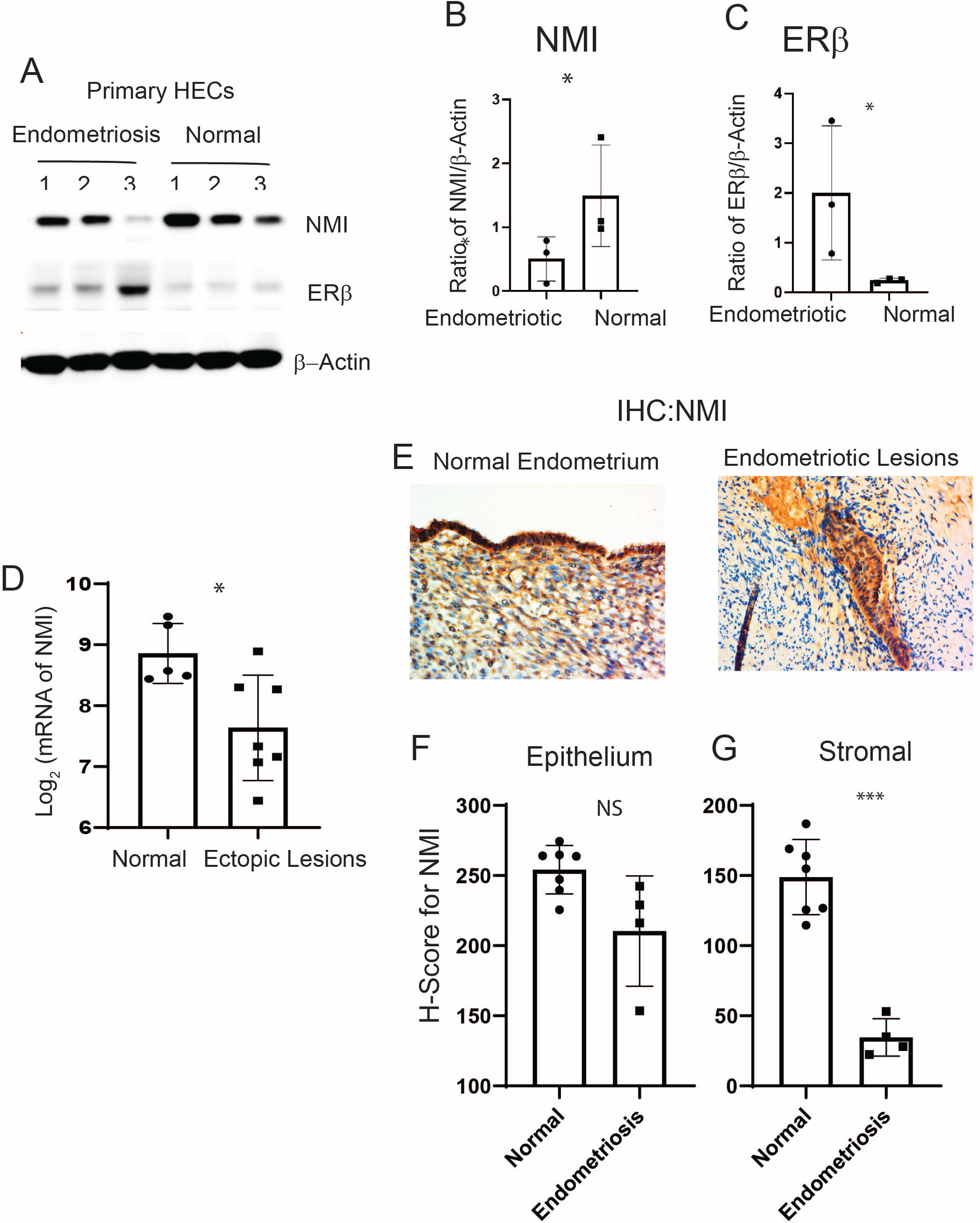
Reduced Levels of NMI in Endometriotic Lesions. (A) Protein levels of NMI, ERβ, and β-Actin in primary human endometrial stromal cells isolated from patients with endometrioma endometriosis (Endometriosis) compared to those from normal endometrium (Normal), as determined by Western blot analysis. (B) The protein ratio of NMI to β-Actin as shown in panel A. (C) The protein ratio of ERβ to β-Actin as shown in panel A. (D) Comparative analysis of NMI mRNA levels in human ectopic lesions versus normal endometrium, based on GSE25628 data. (E) Immunohistochemical analysis of NMI expression in normal human endometrium and human endometriotic lesions. (F) Quantification of NMI protein levels in the epithelial compartment of both the endometrium and endometriotic lesions, as shown in panel E. (G) Quantification of NMI protein levels in the stromal compartment of both the endometrium and endometriotic lesions, as illustrated in panel E. NS: Non-Specific. Significance levels are indicated as follows: *p<0.05, **p<0.01, ***p<0.001.

## The ERβ/HDAC8 axis leads to the reduction of NMI expression in endometrial stromal cells

We generated ERβ-overexpressing endometriotic lesions and eutopic endometrium by auto- transplanting uterine fragments from an endometrium-specific ERβ-overexpressing female mouse back into the same mouse (Han et al., 2015a). For the control group in our endometriosis study, endometriosis was induced by auto-transplanting uterine fragments from control female mice back into the same mouse. Endometriotic lesions and eutopic endometrium were then isolated from both endometrium-specific ERβ-overexpressing mice and their control counterparts with endometriosis. Compared to control ectopic lesions, NMI levels were significantly lower in ectopic lesions from ERβ-overexpressing mice, as evidenced by the NMI ratio in ERβ-overexpressing ectopic lesions to normal ectopic lesions being 0.2 (Fig. 2A) (Han et al., 2019b).

**Fig 2.**
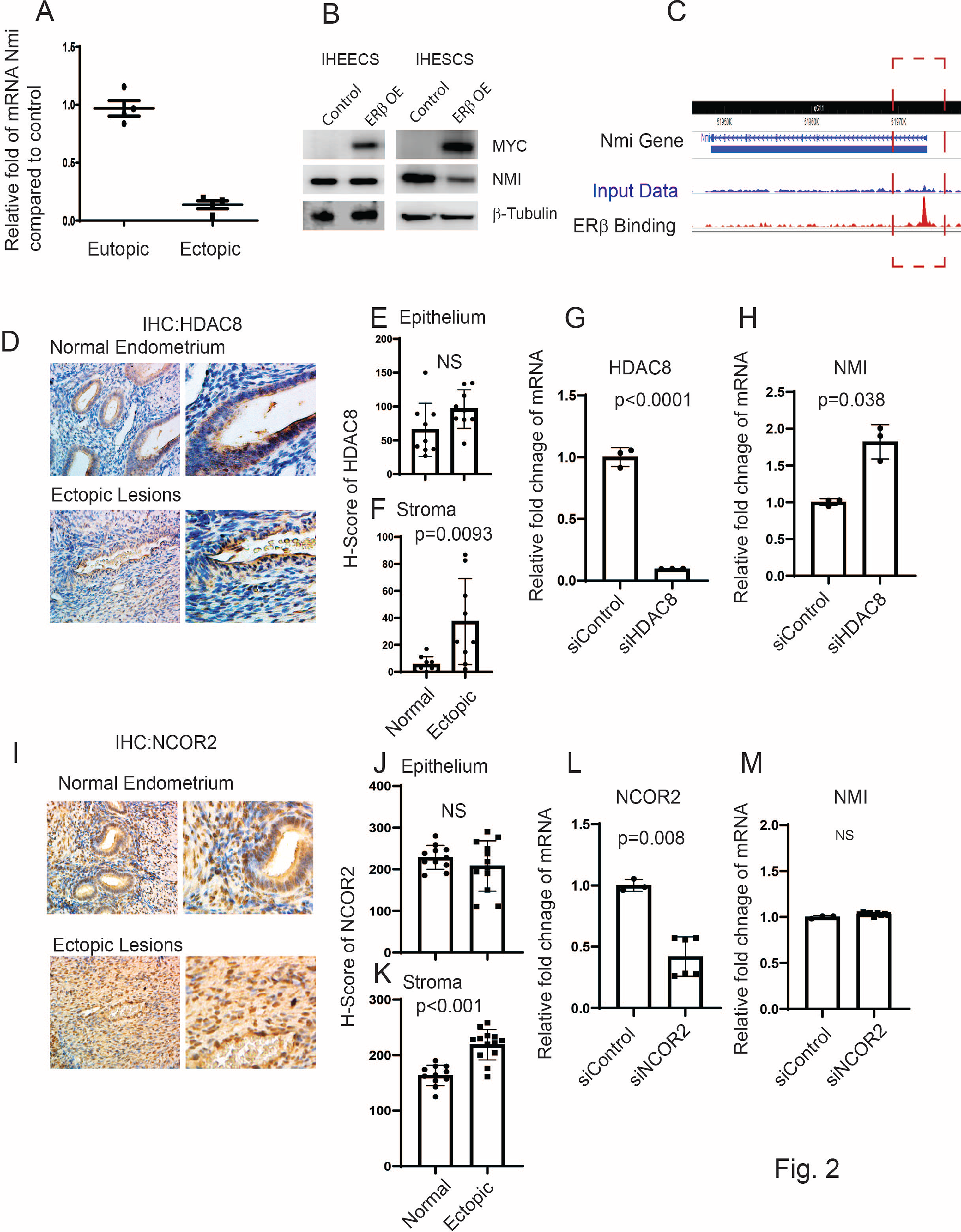
Modulation of NMI Expression via the ERβ/HDAC8 Axis in Endometrial Stromal Cells. (A) Comparative analysis of NMI mRNA levels in ERβ-overexpressing eutopic endometrium and ectopic lesions versus control samples, illustrating the relative fold changes in NMI RNA. (B) Evaluation of NMI protein levels in ERβ-overexpressing immortalized human endometrial epithelial cells (IHEECs) and stromal cells (IHESCs) via Western blot. β-Tubulin served as a normalization control for NMI expression levels, while MYC-tagged ERβ levels were assessed using a MYC antibody. (C) ChIP-seq analysis in ERβ-overexpressing mouse ectopic lesions highlighted ERβ’s binding to the NMI locus promoter region, indicating direct regulatory actions. (D) HDAC8 protein expression in normal endometrium and ovarian endometriotic lesions was analyzed through immunohistochemistry using an HDAC8-specific antibody. (E-F) Quantitative assessment of HDAC8 protein concentrations within the epithelial (E) and stromal (F) compartments of normal endometrium and ovarian endometriotic lesions, as shown in panel D. (G) Quantitative RT-PCR was employed to measure HDAC8 mRNA levels in IHESCs treated with either control siRNA (siControl) or HDAC8-targeting siRNA (siHDAC8). (H) NMI mRNA levels in IHESCs subjected to siControl or siHDAC8 treatments were quantified using quantitative RT-PCR. (I) NCOR2 protein expression in normal endometrium and ovarian endometriotic lesions was determined via immunohistochemistry, utilizing an NCOR2 antibody. (J-K) Quantitative analysis of NCOR2 protein levels in the epithelial (J) and stromal (K) compartments of normal endometrium and ovarian endometriotic lesions, corresponding to panel I. (L) Quantitative RT-PCR was used to ascertain NCOR2 mRNA levels in IHESCs treated with siControl or NCOR2-specific siRNA (siNCOR2). (M) Assessment of NMI mRNA levels in IHESCs following treatment with siControl or siNCOR2, determined by quantitative RT-PCR. NS: Non-Specific.

However, no significant difference in NMI levels was observed between ERβ-overexpressing and wild-type (WT) control eutopic endometrium (Fig. 2A). To further validate our findings, we employed Myc-tagged ERβ-overexpressing immortalized human endometrial epithelial cells (IHEEC) and immortalized human endometrial stromal cells (IHESC) lines that we had previously developed and validated (Han et al., 2015a). MYC-tagged ERβ was robustly expressed in both cell types, as confirmed through Western blot analysis using an MYC-antibody (Fig. 2B). Overexpression of ERβ significantly decreased NMI expression in immortalized human endometrial stromal cells (IHESC) but not in immortalized human endometrial epithelial cells (IHEEC) when compared to their respective parental cells. This pattern aligns with the stromal cell-specific reduction of NMI levels observed in human endometriotic lesions (Fig. 1E), indicating a reverse correlation between ERβ and NMI in endometriotic lesions. The subsequent inquiry focuses on how ERβ reduces NMI levels in endometriotic lesions. To unravel this crucial question, we investigated the potential interaction between ERβ and the promoter regions of NMI genomic loci within endometriotic lesions, utilizing our lesion-specific ERβ ChIP-seq dataset (Han et al., 2019a). ERβ ChIP-seq analysis revealed direct binding of ERβ to the NMI promoter region within endometriotic lesions (Fig. 2C). This raises the question: how does ERβ downregulate NMI levels in human endometrial stromal cells? Several corepressors have been identified that interact with ERβ, thereby inhibiting its activity. (Duong, Licznar et al., 2006, Klinge, Jernigan et al., 2004). Among them, HDAC8 is aberrantly expressed in endometriosis compared to normal tissues (Zheng, Liu et al., 2023a). Immunohistochemical (IHC) analysis comparing human endometriotic lesions with normal endometrium revealed that HDAC8 levels were increased in the stromal cells, but not in the epithelial cells, of human ectopic lesions relative to the normal endometrium (Fig. 2D-F). To investigate the role of HDAC8 in modulating NMI expression in IHESCs, we utilized HDAC8 siRNA to generate HDAC8 knockdown IHESCs (Fig. 2G). Non-target siRNA served as the control. HDAC8 siRNA significantly decreased HDAC8 RNA levels in IHESCs compared to the control siRNA (Fig. 2G). Knocking down HDAC8 using HDAC8 siRNA resulted in an increased RNA level of NMI in IHESCs compared to cells treated with non-target siRNA (Fig. 2H). This indicates that the elevation of HDAC8 in the stroma is associated with decreased NMI levels in endometriotic lesions.

Similarly, the level of NCOR2 was higher in the stromal cells but not in the epithelial cells of human ectopic lesions (Figs. 2I-K). To explore NCOR2’s influence on NMI expression, we reduced the RNA levels of NCOR2 in IHESCs using NCOR2 siRNA, with a control siRNA for comparison (Fig. 2L). However, decreasing NCOR2 did not result in elevated NMI RNA levels in IHESCs compared to the control (Fig. 2M). Thus, unlike HDAC8, the elevation of NCOR2 does not play a role in suppressing NMI levels in the stroma of endometriotic lesions.

### Reducing NMI levels induced an anti-apoptotic phenotype in human endometrial stromal cells following IFNA treatment

IFNA levels are elevated in patients with endometriosis (Park & Han, 2022), and the elevated IFNA causes apoptosis in various cells, including cancer cells (Herzer et al., 2009, Shi et al., 2022). Compared to normal endometrium, a distinctive molecular phenotype of endometriotic cells is their resistance to apoptosis in response to cell-death signaling, including IFNA (Sugihara, Kobayashi et al., 2014). To evaluate if the down-regulation of NMI imparts anti-apoptotic properties in human endometrial stromal cells, countering IFNA-induced cell death signaling, we generated stable NMI-knockdown (KD) IHESC lines (NMI KD-IHESCs) through the introduction of lentiviruses carrying an NMI shRNA expression unit. For control purposes, IHESCs were transduced with a lentivirus containing a non-targeting (NT) shRNA expression unit. Compared to the NT shRNA transduced cells, NMI protein levels were significantly reduced in the NMI KD-IHESCs (Fig. 3A). Subsequently, we investigated the impact of NMI knockdown (KD) on IFNA-induced cell death in IHESCs. Treatment with IFNA significantly inhibited the growth of IHESCs compared to vehicle control (Fig. 3B). In contrast, IFNA treatment did not suppress the growth of NMI KD-IHESCs (Fig. 3B). Thus, NMI KD conferred resistance to apoptosis in IHESCs upon exposure to IFNA.

**Fig 3.**
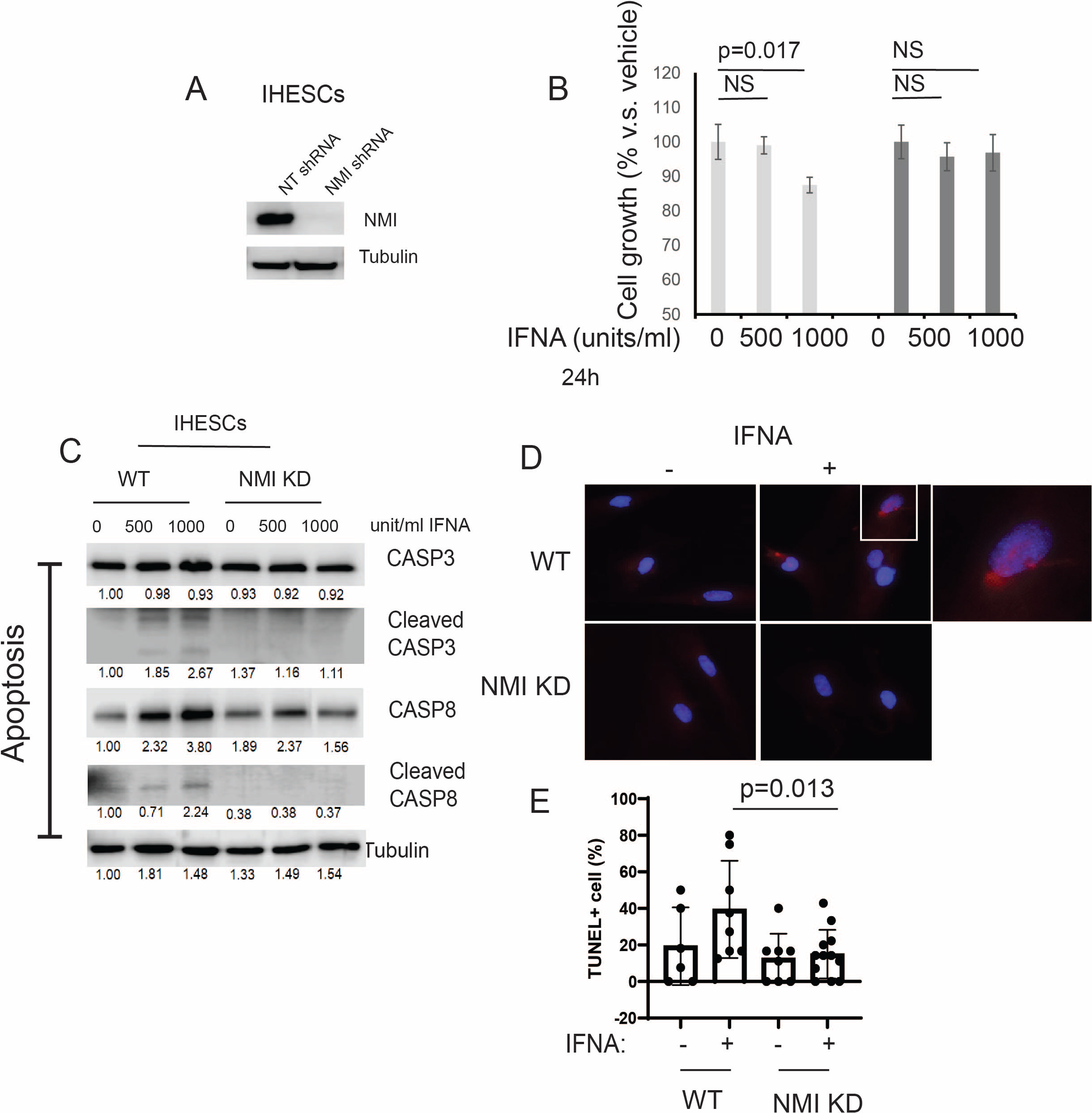
Suppression of IFNα-Induced Apoptosis in Human Endometrial Stromal Cells by NMI Knockdown (KD). (A) Assessment of NMI protein levels in IHESCs treated with non-targeting shRNA (NT shRNA) and NMI-specific shRNA, analyzed via Western blot using an NMI antibody. β-Tubulin was employed as a normalization standard for NMI protein expression. (B) Evaluation of cell proliferation in control IHESCs and NMI KD-IHESCs following treatment with 0, 500, or 1000 units/ml of IFNA for 24 hours, indicating the impact of NMI KD on cell growth under IFNα exposure. (C) Analysis of apoptosis signaling pathways in control KD (wild-type, WT) and NMI KD IHESCs post-treatment with 0, 500, or 1000 units/ml of IFNA for 24 hours. Protein levels of caspase-3 (CASP3), cleaved CASP3, caspase-8 (CASP8), and cleaved CASP8 were quantified via Western blot. β-Tubulin levels served as a normalization control for the apoptosis-associated proteins. (D) Representative images of TUNEL assay in WT and NMI KD IHESCs post- treatment with 0 or 500 units/ml of IFNA for 24 hours. TUNEL positive cells and nuclei were stained in red and blue respectively. Scale bar represents 100µm. (E) Quantitative analysis of TUNEL positivity from (D).

Our previous study demonstrated that ERβ inhibits IFNA-mediated apoptosis in human endometrial epithelial cells by interacting with the apoptosis machinery (Han et al., 2019a). To determine if the loss of NMI also contributes to the prevention of IFNA-induced apoptosis, we exposed both IHESCs and NMI KD-IHESCs to 0, 500, and 1000 units/ml of human recombinant IFNA proteins and analyzed apoptosis signaling through Western blot analysis (Fig. 3C). Treatment with IFNA heightened apoptosis signaling in IHESCs, as evidenced by increased levels of the cleaved forms of CASP3 and CASP8 following IFNA stimulation (Fig. 3C).

Conversely, in NMI KD-IHESCs, NMI knockdown thwarted IFNA-induced apoptosis, indicated by the absence of cleaved CASP3 and CASP8 after IFNA treatment (Fig. 3C). In addition to western blot, TUNEL assay also showed that NMI KD significantly inhibited TUNEL positivity upon IFNA treatment compared to control IHESCs (Fig. 3D, E). Consequently, NMI KD curtailed the IFNA-induced apoptosis signaling in IHESCs.

### NMI KD augmented the expression of genes involved in extracellular matrix signaling and cell adhesion in human endometrial stromal cells

To explore NMI’s involvement in endometriosis progression in depth, we examined the RNA expression profiles of NMI knockdown (NMI KD) in IHESC cells and their respective controls. These cells underwent treatment with either a vehicle or IFNA (1000 unit/ml) for a duration of 24 hours, followed by RNA sequencing. The total read counts for each library, along with the distribution of the transformed data, were illustrated in a bar plot, revealing slight variations in library sizes (Fig. 4A, B). The RNA sequencing process was meticulously performed for each sample. Through hierarchical clustering and Principal Component Analysis (PCA), we identified significant differences in the expression of thousands of genes affected by NMI knockdown in IHESCs. These differences were particularly pronounced when comparing the impact of vehicle and IFNA treatments (Fig. 4C, D). Subsequently, we investigated the molecular phenotype alterations in IHESC cells due to NMI knockdown (NMI KD) under both the absence and presence of IFNA treatment. Initially, this investigation involved analyzing the RNA expression profiles of NMI KD IHESCs versus WT control IHESCs treated with the vehicle. This step aimed to elucidate NMI KD’s impact on IHESCs without IFNA induction. RNA-seq analysis revealed a substantial decrease in NMI RNA levels in NMI KD IHESCs compared to WT controls, with log2 fold changes in NMI RNA levels of -2.98 and -2.99 observed in the presence of both vehicle and IFNA, respectively (Fig. 4E). This outcome confirmed the effective knockdown status of NMI in IHESCs used for RNA sequencing. We then utilized RNA-seq data to pinpoint genes significantly upregulated or downregulated in NMI KD IHESCs relative to WT control cells, adopting thresholds of > 1 (-log10[FDR]) and > 2 (log2[Fold Change]). These findings are depicted in Fig. 4F (Dataset EV1). Further analysis employing Gene Ontology (GO) Biological Process terms highlighted a significant enhancement in extracellular matrix signaling and cell-to-cell adhesion signaling pathways in NMI KD IHESCs compared to WT controls, notably in the absence of IFNA stimulation (Fig. 4G, Dataset EV2).

**Fig 4.**
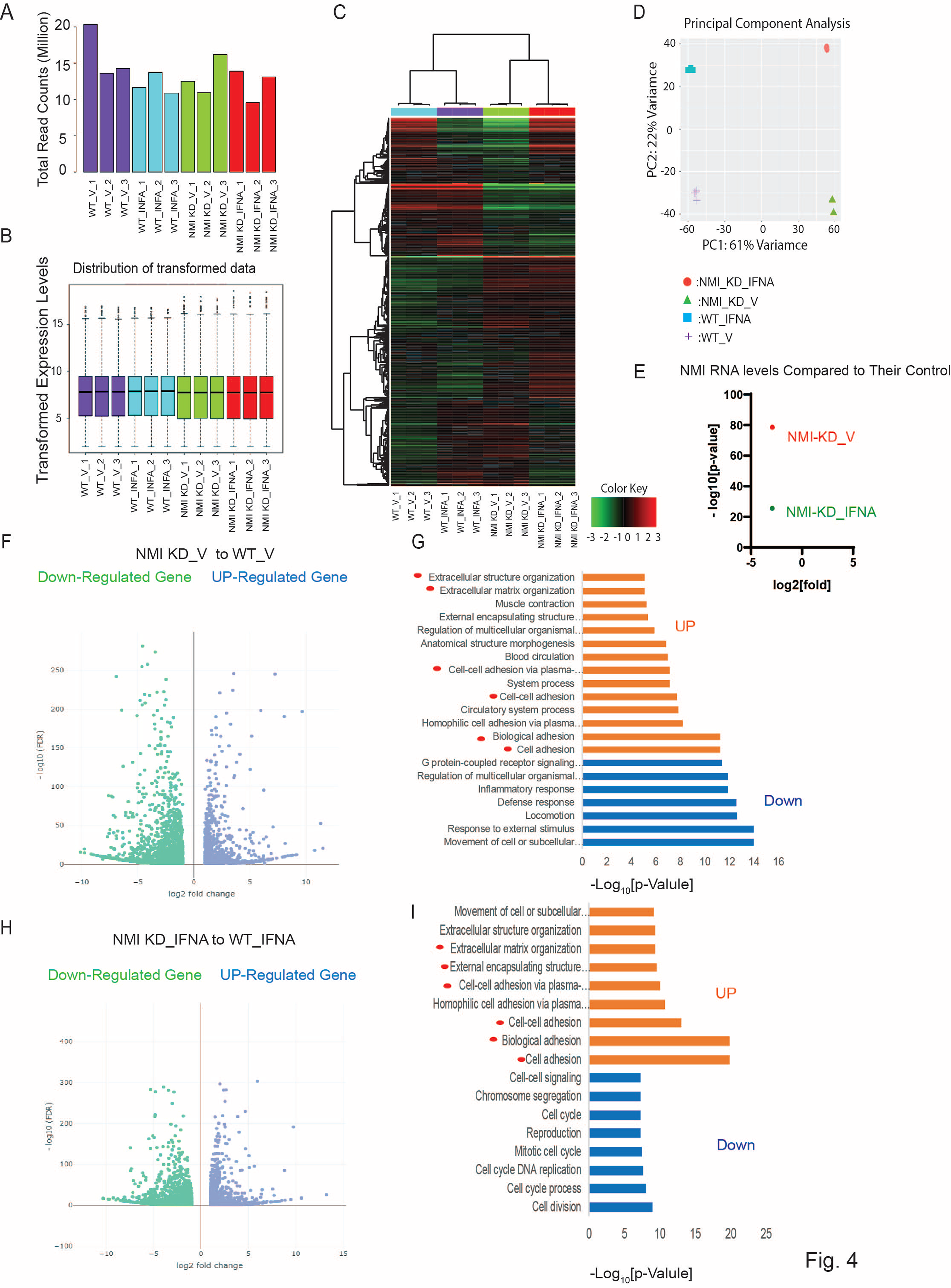
Enhancement of extracellular matrix and cell adhesion signaling in human endometrial stromal cells by NMI KD. (A) Total read counts of each library. (B) Boxplot representing transformed expression levels. (C) Hierarchical clustering and (D) PCA analysis depicting substantial differences in thousands of genes induced by NMI knockdown in IHESCs upon IFNΑ treatment. (E) Relative fold change of NMI RNA levels in NMI KD IHESCs as compared to control KD IHESCs upon vehicle (NMI-KD_V) and IFNΑ (NMI-KD_INFA) treatment. (F) Volcano plot to define the differential gene expression profile between NMI KD and control KD (WT) IHESCs under vehicle treatment. Identification of up- or down-regulated genes with > 1 (-log10[FDR]) and > 2 (log2[Fold]) changes in NMI KD IHESCs compared to WT control IHESCs under vehicle treatment. (G) Gene Ontology analysis heightened the cellular pathways linked to the up-and down-regulated genes in NMI KD IHESCs as compared with control KD IHESCs in Panel F. (H) Volcano plot showing differential gene expression profile between NMI KD and control KD (WT) control IHESCs under IFNA treatment (500 units/ml) for 24 hours. Identification of up- or down-regulated genes with > 1 (-log10[FDR]) and > 2 (log2[Fold]) changes in NMI KD IHESCs compared to WT control IHESCs under IFNA treatment. (I) Gene Ontology analysis heightened the cellular pathways linked to the up-and down-regulated genes in Panel F. (I) Gene Ontology analysis heightened the cellular pathways linked to the up-and down-regulated genes in NMI KD IHESCs as compared with control KD IHESCs upon IFNA treatment in Panel H.

Continuing from there, we assessed the modifications in the RNA expression profile of IHESCs following treatment with IFNA in the context of NMI knockdown. Once again, we identified genes exhibiting significant up- and down-regulation based on the same criteria, comparing NMI KD IHESCs to WT controls following IFNA treatment (Fig. 4H, Dataset EV3). Interestingly, the GO Biological Process analysis unveiled a parallel trend; extracellular matrix and cell-to-cell adhesion signaling pathways were notably enhanced in NMI KD IHSECs compared to WT controls, mirroring the outcomes observed following vehicle treatment (Fig. 4I, Dataset EV4).

To validate RNA sequencing data, the genes MPZL2, ITGB3, JUP, and SORBS1, which are associated with cell adhesion, were chosen to investigate the impact of NMI KD on cell adhesion pathways in endometrial stromal cells. We assessed their mRNA levels in NMI KD and WT control IHESCs following treatment with either vehicle or IFNA using quantitative RT-PCR. NMI KD resulted in elevated mRNA levels of these genes in IHESCs under both conditions, with and without IFNA treatment, compared to the WT control (Fig. 5A, B). Treatment with IFNA significantly increased the RNA levels of these genes in NMI KD-IHESCs in comparison to the vehicle treatment. Additionally, the mRNA levels of MFAP5, ADAM19, VCAN, and HAPLN3, which are implicated in extracellular matrix signaling, were evaluated in NMI KD and WT control IHESCs following treatment with vehicle or IFNA, using quantitative RT-PCR. NMI KD led to an increase in mRNA levels of all these genes involved in extracellular matrix signaling in IHESCs, in both the absence and presence of IFNA, when compared to the WT control (Fig. 5C, D). Treatment with IFNA significantly enhanced the expression of these genes in NMI KD-IHESCs relative to the vehicle. In summary, the downregulation of NMI in IHESCs augmented signaling pathways associated with the extracellular matrix and cell adhesion, both crucial for the advancement of endometriosis in response to IFNA exposure.

**Fig 5.**
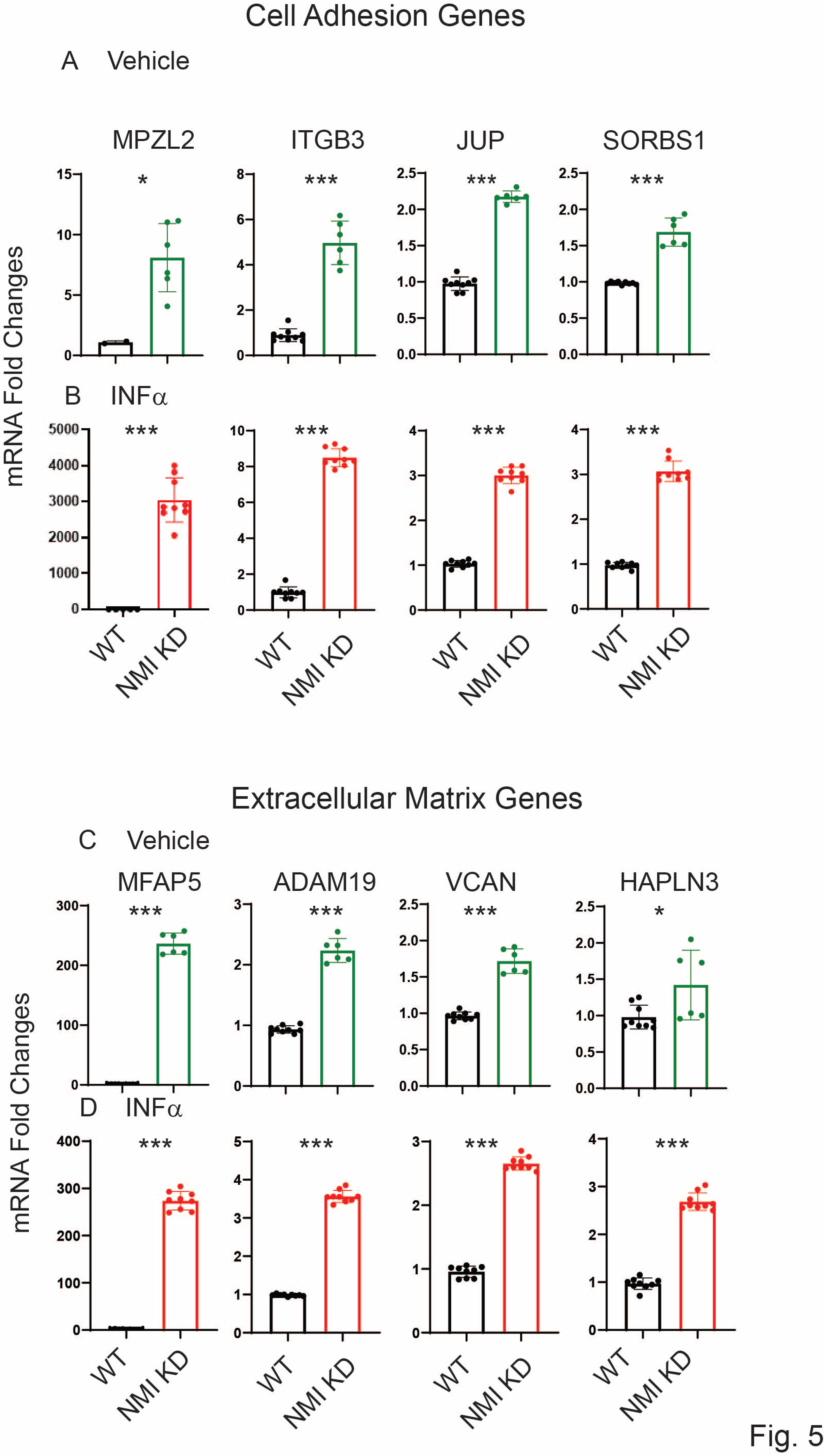
Upregulation of Genes involved in Cell Adhesion and Extracellular Matrix Signaling in NMI KD IHESCs compared to control KD IHESCs. (A) mRNA fold changes of cell adhesion-related genes (MPZL2, ITGB3, JUP, and SORBS1) in control KD (WT) versus NMI KD IHESCs with vehicle treatment and (B) IFNA treatment at 500 units/ml for 24 hours. (C) mRNA fold changes of genes associated with extracellular matrix signaling (MFAP5, ADAM19, VCAN, HAPLN3) in WT versus NMI KD IHESCs with vehicle treatment and (B) IFNA treatment at 500 units/ml for 24 hours. Significance levels are indicated as follows: *p<0.05, **p<0.01, ***p<0.001.

### NMI KD inhibited the apoptosis and necroptosis mediated by interferon-stimulated genes (ISGs) in IHESCs upon stimulation with IFNA, affecting the IFNA-regulated canonical pathway

Exposure to IFNA stimulates the formation of the Interferon-stimulated gene factor 3 (ISGF3) complex, composed of phosphorylated STAT1 (P-STAT1), phosphorylated STAT2 (P-STAT2), and Interferon Regulatory Factor 9 (IRF9) and then ISGF3 complex binds to the Interferon-sensitive response element (ISRE) then enhances g the expression of ISGs, including those involved in cell death signaling (Mazewski et al., 2020, Sarhan, Liu et al., 2019). Given the crucial role of the ISGF3 complex in IFNA signaling pathways (McComb, Cessford et al., 2014), we postulated that knocking down NMI might hinder the formation of the IFNA-induced ISGF3 complex, thereby preventing IFNA-induced cell death signaling in endometriotic cells.

IFNA treatment led to a significant increase in the total protein expressions of IRF9, STAT1, phosphorylated STAT1 (p-STAT1), STAT2, and phosphorylated STAT2 (p-STAT2) in IHESCs, as opposed to vehicle-treated cells (Fig. 6A). Contrary to protein levels, RNA-seq data indicated that NMI knockdown (NMI-KD) did not alter the RNA levels of IRF9 and STAT1/2 in IHESCs compared to control (Fig. 6B). Hence, IFNA treatment notably enhanced the formation of the ISGF3 complex in IHESCs. However, NMI knockdown resulted in reduced protein levels of IRF9, STAT1, and STAT2 in IHESCs upon IFNA stimulation, without a corresponding decrease in their RNA levels (Figs. 6A, B). Additionally, NMI KD diminished the activation of STAT1 and STAT2, as determined by the ratios of p-STAT1/STAT1 and p-STAT2/STAT2, in IHESCs upon IFNA stimulation compared to control IHESCs (Figs. 6C, D). Therefore, NMI may play a role in stabilizing the protein levels of IRF9, STAT1, and STAT2 upon IFNA stimulation, thus facilitating the formation of the ISGF3 complex and elevating ISGs. These genes are implicated in the progression of apoptosis by regulating apoptosis-associated genes (Chawla-Sarkar, Lindner et al., 2003). RNA-seq analysis revealed that NMI knockdown (NMI KD) significantly decreased mRNA levels of apoptotic genes, including Death-Associated Protein Kinase 1 (DAPK1), DAPK2, Caspase 8 (CASP8), Caspase 8 Associated Protein 2 (CASP8AP2), Interferon Regulatory Factor 1 (IRF1), and IRF6 in IHESCs, in comparison to control IHESCs following IFNA stimulation (Fig. 6E). Consequently, NMI KD markedly attenuated IFNA-induced apoptosis in IHESCs. This specific effect of NMI knockdown, which disrupts ISGF3 complex formation, contributes to the anti-apoptotic property of endometriotic lesions, countering cell-death signaling such as IFNA-induced apoptosis.

**Fig 6.**
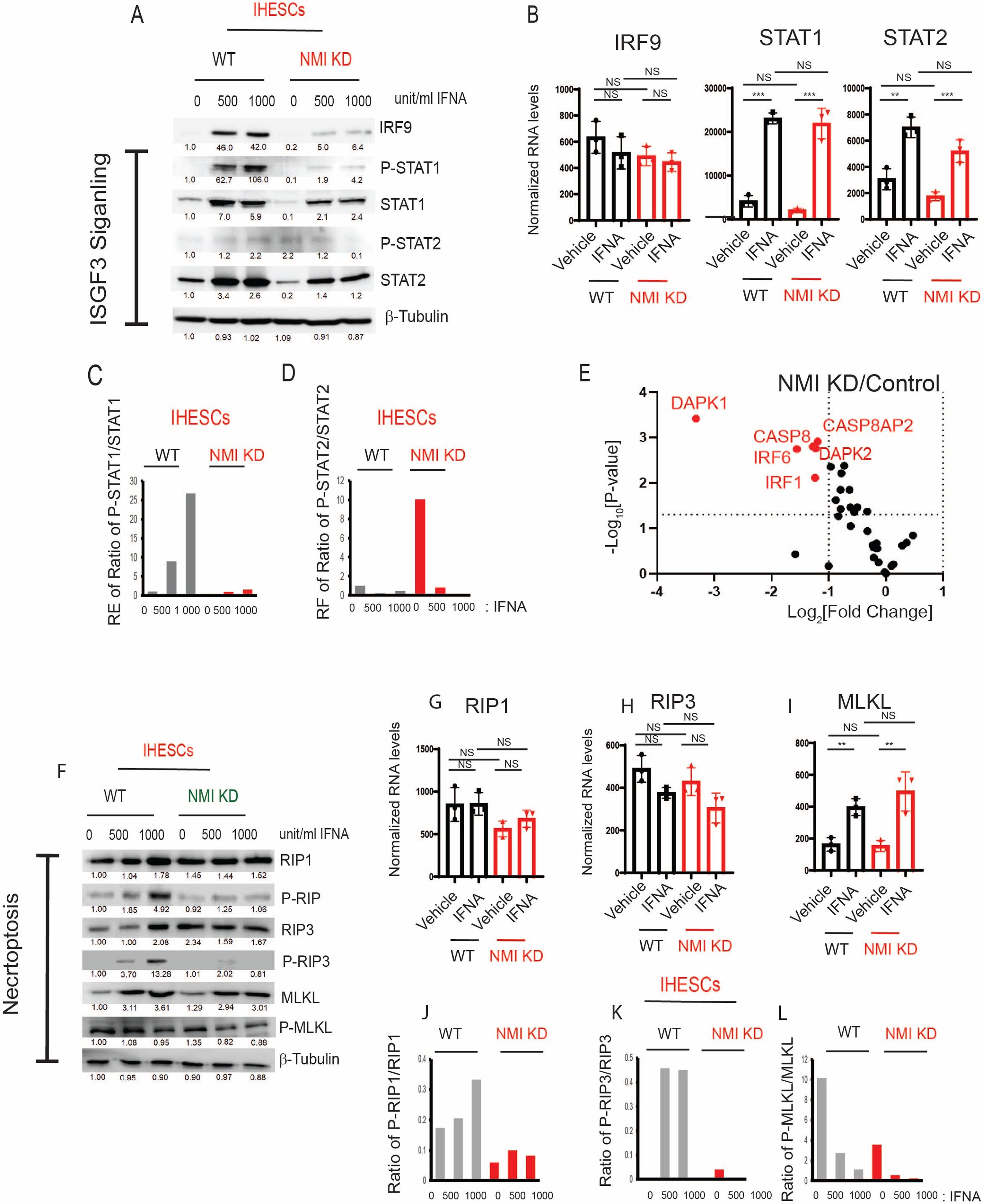
Suppression of ISGF3-mediated apoptosis and Necroptosis in human endometrial stromal cells by NMI KD upon IFNA treatment. (A) Protein levels of ISGF3 complex components, IRF9, STAT1, p-STAT1(Tyr701), STAT2, p-STAT2 (Tyr689) in control KD versus NMI KD IHESCs upon IFNA treatment determined by Western blotting. (B) RNA levels of IRF9, STAT1, and STAT3 normalized by 18sRNA in control KD versus NMI KD IHESCs upon IFNA treatment. (C) The ratio of p-STAT1/STAT1 and (D) the ratio of p-STAT2/STAT2 in control KD versus NMI KD IHESCs upon IFNA treatment determined by quantification of panel A. (E) The relative fold change of RNA levels of genes involved in apoptosis in NMI KD IHESCs compared to control KD IHESCs upon IFNA treatment. (F) Protein levels of necroptosis components. RIP1, p-RIP1(Ser166), RIP3, p-RIP3(Ser227), MLKL, and p-MLKL(Ser358) in control KD (WT) versus NMI KD IHESCs following IFNA treatment. RNA levels of (G) RIP1, (H) IRIP3, and (I) MLKL normalized by 18sRNA in control KD versus NMI KD IHESCs upon IFNA treatment. The ratio of (J) p-RIP1/RIP1, (K) p-RIP2/RIP2, and (L) p-MLKL/MLKL in control KD versus NMI KD IHESCs upon IFNA treatment determined by quantification of panel F. NS: Non-Specific. Significance levels are indicated as follows: *p<0.05, **p<0.01, ***p<0.001.

Necroptosis, an alternative mode of cell death, is also regulated by IFN via the activation of Receptor-Interacting Protein Kinase 1 (RIP1) and Receptor-Interacting Protein Kinase 3 (RIP3) (Vanden Berghe, Linkermann et al., 2014). Type I IFN initiates the phosphorylation of RIP1 and RIP3, which subsequently recruits and activates Mixed Lineage Kinase Domain-Like (MLKL) to assemble the necrosome, thereby inducing cell death (Vanden Berghe et al., 2014). In macrophages, IFN induces necroptosis through ISGF3 signaling by persistently activating RIP3, ultimately leading to inflammation (McComb et al., 2014). Given that NMI knockdown adversely affected ISGF3 signaling in endometrial stromal cells, we explored whether NMI knockdown could also inhibit necroptosis in these cells, potentially facilitating the progression of endometriosis. To assess necroptosis, we examined the protein levels of RIP1, RIP3, and MLKL in IHESCs. NMI knockdown did not affect the protein levels of these necroptosis markers in IHESCs following IFNA treatment (Fig. 6F). Similarly, NMI knockdown did not alter the mRNA levels of RIP1, RIP3, and MLKL in IHESCs upon IFNA treatment (Figs. 6G-I).

Subsequently, we assessed the activation of RIP1, RIP3, and MLKL by examining their phosphorylated forms.

We calculated the ratios of phosphorylated RIP1 (Ser166) (p-RIP1) to RIP1, phosphorylated RIP3 (Ser227) (p-RIP3) to RIP3, and phosphorylated MLKL (Ser358) (p-MLKL) to MLKL based on Western blot data (Fig. 6F). IFNA treatment significantly increased the ratios of p- RIP1/RIP1 and p-RIP3/RIP3 in IHESCs compared to vehicle-treated cells (Fig. 6J, K). However, NMI knockdown suppressed the elevation of the p-RIP1/RIP1 and p-RIP3/RIP3 ratios in IHESCs upon IFNA stimulation compared to vehicle (Fig. 6J, K), thereby preventing the activation of RIP1 and RIP3 in IHESCs upon IFNA stimulation. Interestingly, IFNA treatment decreased the ratio of p-MLKL/MLKL in IHESCs compared to vehicle-treated cells (Fig. 6L). NMI knockdown significantly reduced the ratio of p-MLKL/MLKL in IHESCs upon IFNA stimulation compared to control IHESCs (Fig. 6L). Therefore, NMI knockdown effectively suppressed the phosphorylation of necroptosis components upon IFNA treatment and hindered IFNA-induced necroptosis in endometrial stromal cells, potentially contributing to the progression of endometriosis.

### NMI knockdown enhanced IFNA-induced activation of non-canonical pathways in human endometrial stromal cells

While the JAK/STAT pathways are recognized as canonical pathways activated by IFNA, this cytokine also triggers various non-canonical pathways, including PI3K/AKT, GSK3β/β-catenin, and MAPK, to modulate cellular processes (Li, 2008, Marineau, Khan et al., 2020, Mazewski et al., 2020). Furthermore, the activation of these non-canonical IFNA pathways plays a crucial role in the progression of endometriosis (Madanes, Bilotas et al., 2020, Pazhohan, Amidi et al., 2018, Santulli, Marcellin et al., 2015). For example, the aberrant activation of Wnt/β-catenin signaling during the secretory phase of the menstrual cycle in endometriosis has been associated with excessive inactivation of GSK3β (Pazhohan et al., 2018). Furthermore, NMI inhibits AKT and GSK3β/β-catenin signaling in various cancers, thereby impeding cancer progression (Fillmore et al., 2009, Wang et al., 2017). However, the biological functions of NMI in IFNA-induced non-canonical cellular pathways during the progression of endometriosis remain unexplored. To investigate this, we aimed to determine the effects of NMI knockdown on GSK3β/β-catenin and AKT signaling in endometrial stromal cells. Following IFNA treatment, there was no change in the protein levels of β-catenin, but GSK3β levels were increased in IHESCs (Fig. 7A).

**Fig 7.**
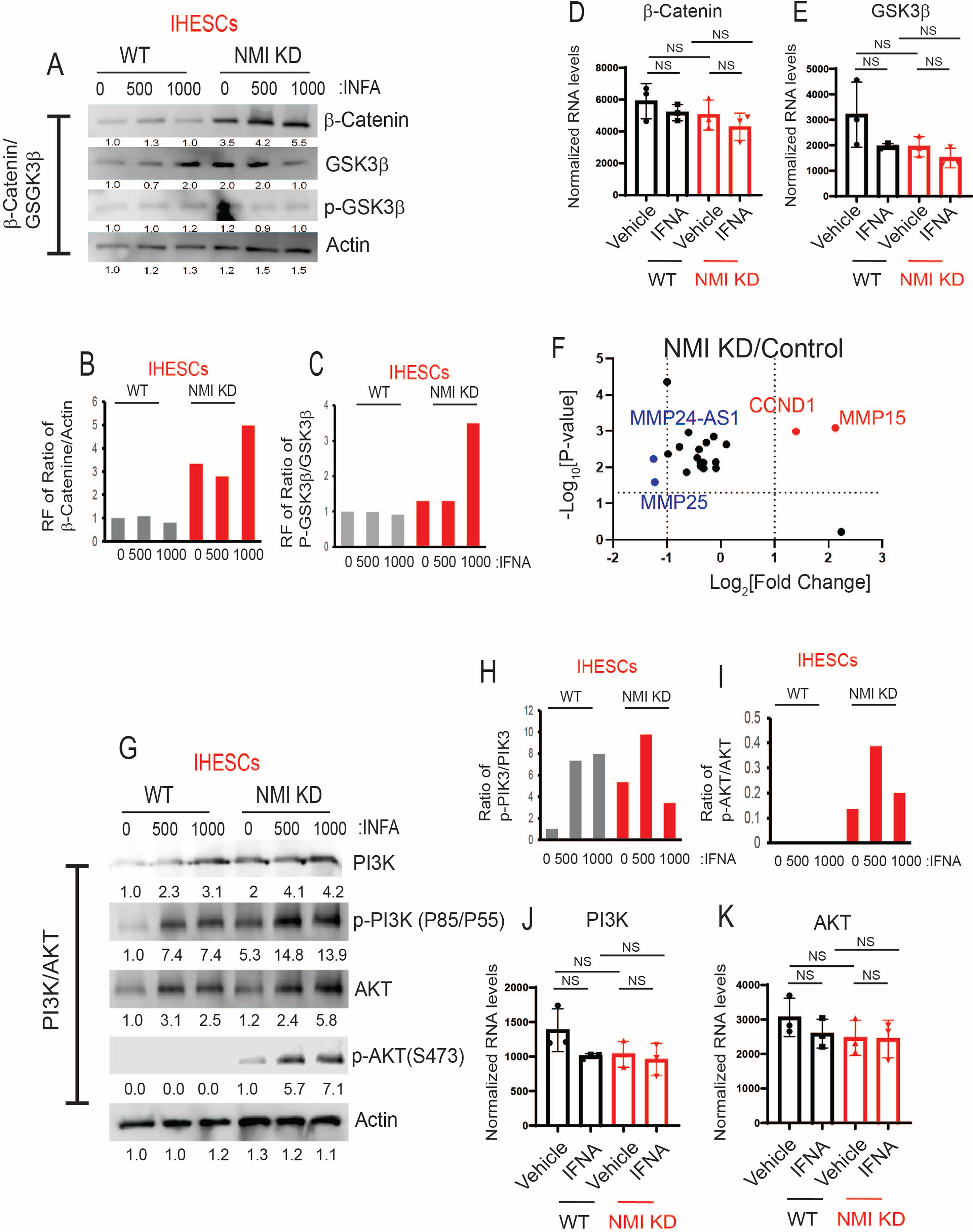
Increasing IFNA Non-Canonical Pathways in Endometrial Stromal Cells by NMI KD. (A) Protein levels of β-Catenin, GSK3β, and p-GSK3β (Ser9) in control KD (WT) versus NMI KD IHESCs upon IFNA treatment. Protein levels of b-actin were determined to normalize the β-Catenin/GSK3β axis. RNA levels of (B) β-Catenin and (C) GSK3β normalized by 18SRNA were determined in control KD (WT) and NMI KD IHESCS upon IFNA treatment. Relative fold change of (D) the ratio of β-Catenin/b-Actin and (E) the ratio of p-GSK3β/GSK3β in control KD (WT) and NMI KD IHESCS upon IFNA treatment were calculated based on panel A. (F) The relative fold change of RNA levels of β-Catenin target genes in NMI KD IHESCs compared to control KD IHESCs. (G) Protein levels of PI3K, p-PI3K (Tyr 85/Tyr 55), AKT, and p-AKT (Ser473) in control KD (WT) and NMI KD IHESCs upon IFNA treatment. Protein levels of b-actin were determined to normalize the PI3K/p-PI3K and the AKT/p-AKT axis. (H) Relative fold change of the ratio of p-PI3K(Tyr85/55)/PI3K in control KD (WT) and NMI KD IHESCs upon IFNA treatment was quantified based on panel G (I) Relative fold change of the ratio of p-AKT(Ser473)/AKT in control KD (WT) and NMI KD IHESCS upon IFNA treatment were calculated based on panel G. (J) RNA levels of PI3K normalized by 18sRNA were determined in control KD (WT) and NMI KD IHESCS upon IFNA treatment (K) RNA levels of AKT normalized by 18sRNA were determined in control KD (WT) and NMI KD IHESCS upon IFNA treatment. NS: Non-Specific.

Meanwhile, the RNA levels of β-Catenin and GSK3β remained unchanged in IHESCs upon IFNA treatment (Figs. 7B, C). Compared to the control, NMI knockdown (KD) significantly increased β-Catenin protein levels in immortalized human endometrial stromal cells (IHESCs), and IFNA treatment further elevated β-Catenin protein levels in NMI KD-IHESCs without affecting its RNA expression (Fig. 7A, B). Conversely, NMI KD led to a decrease in GSK3β protein levels in IHESCs following IFNA treatment, without a corresponding reduction in its RNA expression (Fig. 7A, C). Subsequently, we assessed the inhibition of GSK3β activity through the phosphorylation of serine 9 by Western blot analysis. IFNA treatment alone did not alter the phosphorylated GSK3β (p-GSK3β) to GSK3β ratio in IHESCs (Fig. 7A, E). However, in the context of NMI KD and IFNA treatment, there was a significant increase in the p- GSK3β/GSK3β ratio, indicating a reduction in GSK3β activity in IHESCs (Fig. 7A, E). Further investigation into how NMI KD affects the expression of β-Catenin target genes revealed that RNA sequencing (RNA-seq) analysis demonstrated a significant upregulation of CCND1 and MMP15 RNA levels in IHESCs with NMI KD compared to the control (Fig. 7F). Both CCND1 and MMP15 have an essential role in cell proliferation (Alam, Zunic et al., 2022, Majali- Martinez, Hoch et al., 2020). Therefore, NMI knockdown enhanced the β-catenin/GSK3β axis in endometrial stromal cells, amplifying the effects of IFNA-induced non-canonical pathways, which in turn increased the proliferation of endometriotic lesions.

Subsequently, we explored the effects of NMI knockdown on the IFNA-induced non-canonical PI3K/AKT pathways. In IHESCs, IFNA treatment resulted in an increase in PI3K and AKT protein levels without a corresponding increase in their mRNA levels (Fig. 7G, J, K). NMI KD increased PI3K protein levels, which but did not lead to an increase in its mRNA levels (Fig. 7G, J). IFNA treatment also increased levels of phosphorylated PI3K (p-PI3K) in IHESCs (Fig. 7G, H). However, NMI KD did not further increase activated PI3K induced by IFNA (Fig 7G, H). In addition, NMI KD did not affect the increase in AKT protein levels induced by IFNA treatment, which also did not lead to an increase in mRNA levels in IHESCs (Fig. 7G, K). Despite the elevation of AKT protein levels in IHESCs, IFNA treatment alone did not activate AKT, as indicated by unchanged levels of phosphorylated AKT (p-AKT) at Ser473 (Fig. 7G, I). However, NMI KD resulted in an increase in p-AKT levels in IHESCs following IFNA stimulation (Fig. 7I). Consequently, NMI KD contributed to the activation of PI3K/AKT signaling in IHESCs, as evidenced by a significant increase in the ratio of p-AKT (Ser473) to AKT in NMI KD-IHESCs upon IFNA treatment (Fig. 7G, I).

### NMI KD endometrial stromal cells enhanced the growth of ectopic lesions in mice with endometriosis

NMI knockdown inhibited IFNA-induced cell death and enhanced non-canonical IFNA pathways in endometrial stromal cells. Consequently, we investigated whether suppressing NMI in these cells could promote the growth of ectopic lesions in mice, mirroring human endometriosis. To this end, we peritoneally injected mice with a mixture of luciferase-labeled control wild-type endometrial epithelial cells (LIHEECs) and luciferase-labeled control wild-type endometrial stromal cells (LIHECs) (referred to as Epi+Str), along with control wild-type LIHEECs and NMI KD-LIHESCs (referred to as Epi+NMI KD-Str). Since both cell types were luciferase-labeled, we could non-invasively monitor the growth of the injected cell mixtures via luciferase activity. After injection, the mixtures of endometrial stromal and epithelial cells localized well in the pelvic area of the recipient female mice (Fig. 8A). On the 10th day post-endometriosis induction, we assessed the luciferase activity emanating from the human ectopic lesions formed in NOD-SCID mice. This analysis revealed that the mixture containing epithelial cells with NMI KD-stromal cells (Epi+NMI KD-Str) demonstrated a significantly more pronounced development of endometriotic lesions compared to the control mixtures of epithelial and stromal cells, as indicated by the significantly higher luciferase activity in stromal NMI KD endometriotic lesions compared with control lesions (Fig. 8A, B). In addition to measuring luciferase activity, we also quantified the number of ectopic lesions per mouse. The combination of epithelial cells with NMI KD-stromal cells (Epi+NMI KD-Str) resulted in a higher count of ectopic lesions than the control mixture of endometrial epithelial and stromal cells (Fig. 8A, C). Thus, the suppression of stromal NMI was found to enhance the growth of ectopic lesions in mice models of endometriosis.

**Fig 8.**
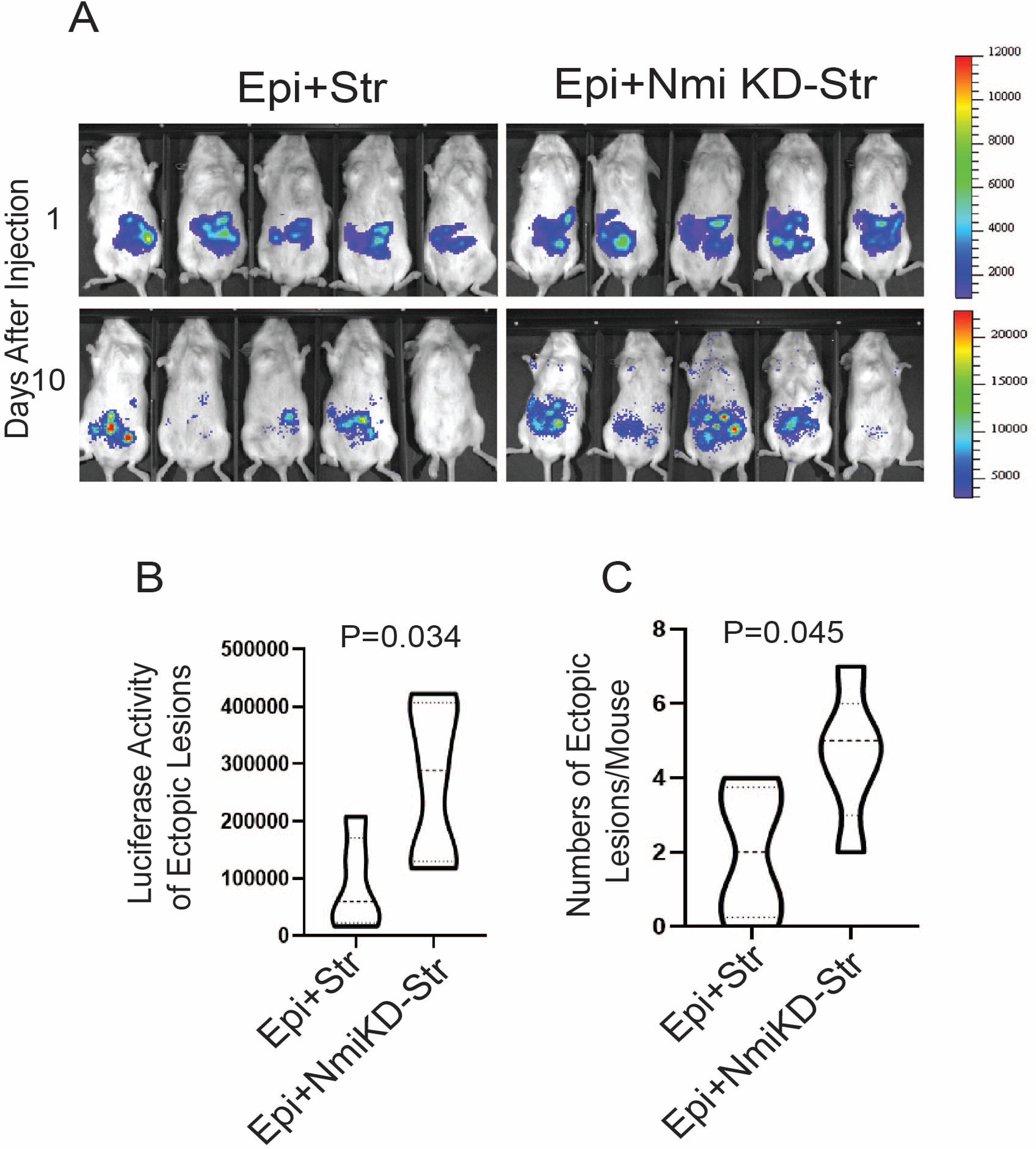
Stimulation of growth of endometriotic lesions in mice by NMI KD endometrial stromal cells. (A) Bioluminescence images of endometriotic lesions generated by the mixture of luciferase labeled immortalized human epithelial cells (LIHEEC) plus luciferase labeled immortalized human stromal cells (LIHESC) named as “Epi+Str”, and mixture of LIHEECS plus NMI KD LIHESCs named as Epi+NMI KD-Str in SCID mice at 0 and 10 days post- endometriosis induction. (B) Quantification of bioluminescence signals depicted in panel A. (C) Quantification of the number of ectopic lesions corresponding to the bioluminescence signals from panel A.

## Discussion

In reproductive-aged women, retrograde menstruation leads to the dispersal of endometrial fragments within the pelvic region. Plasmacytoid dendritic cells (pDCs) recognize and present endometrial autoantigens, thereby increasing type I IFN level (Suen, Chang et al., 2019). This elevation contributes to the differentiation of monocytes, sustains the survival of dendritic cells, and enhances the activity of cytotoxic T cells (Ali, Mann-Nüttel et al., 2019, Newby, Brusko et al., 2017). These processes potentially enhance the elimination of endometrial fragments in the pelvic area, especially in healthy women (Pia et al., 2012). Our findings revealed that exposure to IFNA increased cell death mechanisms, such as apoptosis and necroptosis, in endometrial stromal cells, effectively inhibiting their proliferation. Nonetheless, irregular IFNA signaling is pivotal in the progression of endometriosis, distinguishing affected individuals from healthy women. Notably, transcriptions of IFNA and IFNAR2 are significantly upregulated in the endometrium of women with endometriosis compared to their healthy counterparts (Kao et al., 2003). Furthermore, JAK1, a critical component of type I IFN signaling, shows increased expression in endometriosis stromal cells compared to endometrial cells from individuals without endometriosis (Matsuzaki et al., 2006). A study involving fifty-two infertile patients diagnosed with moderate or severe endometriosis showed that the intraperitoneal administration of human recombinant IFNA-2b following endometriosis surgery resulted in a recurrence of endometriosis after 21 months (Acien, Quereda et al., 2002). These findings suggest an essential role of type I IFN in endometriosis progression. However, a study presents evidence of inhibiting endometriosis progression by IFNA. For instance, the contested treatment involving Human IFNA-2b demonstrated the suppression of proliferation and migration in primary endometrial stromal cells sourced from patients, and this treatment attenuates endometriosis in a rat model (Dicitore, Castiglioni et al., 2018, Ingelmo, Quereda et al., 1999).

Our previous study indicated that ERβ inhibits TNFα-induced apoptosis signaling while simultaneously promoting the proliferation, adhesion, and invasion activities of human endometriotic epithelial cells (Han et al., 2015a). However, it remains unclear how endometriotic stromal cells bypass apoptosis signaling to develop into endometriotic lesions in patients with endometriosis. In this context, we propose that downregulated NMI plays a critical role in the survival of endometrial stromal cells against host immunosurveillance. The downregulation of NMI facilitates the avoidance of IFNA-induced cell death signaling and amplifies IFNA non-canonical pathways within endometrial stromal cells, thereby furthering the progression of endometriosis (Fig. 9). Additionally, apart from endometriosis, NMI also exhibits cancer-suppressive properties in various cancers. For example, NMI facilitates the upregulation of dickkopf 1 (Dkk1), an inhibitor of the Wnt/β-catenin signaling pathway. This regulatory action suppresses Wnt/β-catenin signaling and subsequently limits the growth of breast tumors in vivo (Fillmore et al., 2009). NMI also restricts the growth and migration of lung cancer cells by enhancing cell apoptosis (Wang et al., 2017). Therefore, NMI exhibits suppressive activity against breast and lung cancer, as well as endometriosis, through a similar mechanism. However, it should be noted that NMI demonstrates an oncogenic function in other types of cancer, such as glioblastoma (He, Qiao et al., 2022, Pruitt, Devine et al., 2016). Hence, NMI possesses a tissue-specific function, dynamically modulating cellular pathways.

**Fig 9.**
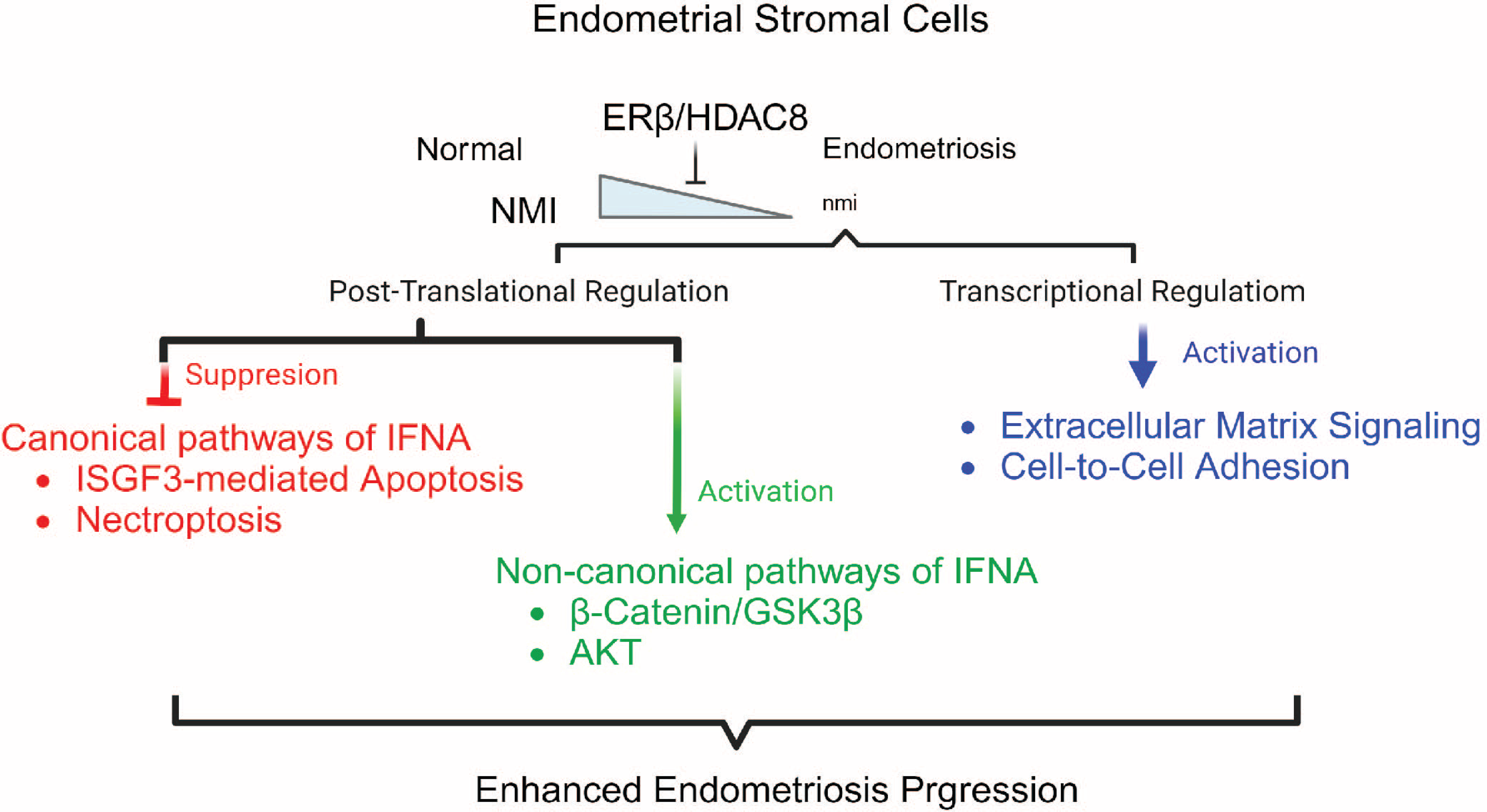
Model for endometriosis progression by downregulating NMI in endometrial stromal cells The activation of the ERβ/HDAC8 axis leads to decreased NMI levels in endometrial stromal cells, which in turn interferes with the formation of the ISGF3 complex by promoting the degradation of IRF9 and STAT1/2 proteins. This process effectively blocks IFNα-induced apoptosis. Additionally, the reduction in NMI levels prevents IFNA-triggered necroptosis by inhibiting the activation of RIP1/RIP2 and MLKL in these cells. Beyond the suppression of canonical IFNα pathways, diminished NMI levels facilitate the activation of non-canonical IFNα signaling routes. For instance, lower NMI levels enhance β-Catenin/GSK3β signaling by stabilizing β-Catenin protein and activating AKT signaling, thereby promoting cell proliferation in response to IFNα treatment. Furthermore, decreased NMI expression upregulates RNA levels of genes associated with extracellular matrix signaling and cell-to-cell adhesion, contributing to the exacerbation of endometriosis.

IHC analysis of human endometriotic lesions revealed a significant reduction in NMI levels within stromal cells. However, no significant decrease was observed in epithelial cells of ectopic lesions compared to healthy women’s normal endometrium. How does NMI lead to stromal-specific reduction in endometriotic lesions? Understanding the role of down-regulated stromal NMI in endometriosis progression is critical. Histone deacetylation typically silences genes, as HDACs function as transcriptional repressors (Chen, Zhao et al., 2015). HDACs play a significant role in endometriosis. For instance, HDAC inhibition resulted in the reactivation of E-cadherin, attenuation of invasion, and decreased proliferation of endometriotic cells (Wu, Starzinski-Powitz et al., 2007). Trichostatin A, an HDAC inhibitor, suppressed endometriotic lesion growth and hyperalgesia in a mouse model of endometriosis (Lu, Nie et al., 2010). A recent study revealed that HDAC8 is progressively and aberrantly overexpressed as endometriotic lesions advance (Zheng et al., 2023a, Zheng, Liu et al., 2023b). However, this study did not elucidate the role of HDAC8 in the progression of endometriosis. Here, we propose that the ERβ/HDAC8 axis is the key driver for the downregulation of endometrial stromal NMI to enhance endometriosis. Additionally, there is a potent HDAC8-specific inhibitor, PCI-34051, with over 200-fold selectivity over other HDAC isoforms (Balasubramanian, Ramos et al., 2008), and the treatment with PCI-34051 potentiates antitumor immunity (Yang, Feng et al., 2021). Our previous study revealed that oleuropein selectively suppressed ERβ compared to ERα in endometriotic lesions (Park et al., 2022). Therefore, the combination of PCI-34051 and oleuropein might effectively suppress the ERβ/HDAC8 axis and consequently restore NMI expression in endometrial stromal cells, rendering them sensitive to IFNA-induced cell death signaling for endometriosis treatment. We will investigate whether the combination of PCI- 34051 and oleuropein effectively suppresses the progression of endometriosis.

NMI interacts with various transcription factors and co-regulators to dynamically regulate gene expression (Pruitt et al., 2016). Therefore, dysregulation of NMI-mediated gene regulation is associated with human disease progression. For example, down-regulation of NMI expression in breast cancer and lung cancers leads to aberrant expression of oncogenes or dysregulation of tumor suppressor genes, contributing to tumorigenesis and cancer progression (He et al., 2022, Wang et al., 2017). Therefore, dysregulation of NMI-mediated transcriptional inhibition can enhance oncogenic cellular pathways in these cancers. In the case of endometriosis, NMI effectively suppresses cell-to-cell adhesion and extracellular matrix signaling to prevent endometriosis progression. Thus, dysregulation of NMI-mediated transcriptional inhibition elevates cell-to-cell adhesion and extracellular matrix signaling to enhance endometriosis progression. However, how NMI inhibits transcription of these genes needs to be addressed. A recent study showed that the interaction of NMI with IFP35 and then inhibits IFP35 protein degradation (Li, Chen et al., 2023). Therefore, NMI directly stabilizes the IFP35 protein through its interaction with IFP35. NMI is known to interact with components of the IRF9/STAT1/STAT2 complex. Consequently, NMI’s integration into ISGF3 complex components might stabilize these proteins, enhancing ISGF3 complex-mediated IFNA-induced apoptosis. However, how NMI increases protein stability through interaction has not been elucidated yet. In addition to protein stabilization, NMI is also involved in protein destabilization. For instance, NMI interacts with IRF3 and IRF7, promoting the autophagy-mediated degradation of these two transcription factors (Li et al., 2023). In endometriosis, NMI also destabilizes β-catenin in normal endometrial stromal cells. Inhibition of GSK3β stabilized β- catenin protein (Wu, Huang et al., 2009). Therefore, the stabilization of β-catenin by NMI knockdown would be generated by inactivating GSK3β through an increase in p-GSK3β (Ser9).

NMI was highly expressed in normal lung cells, but its expression in lung cancer cells was significantly reduced (Wang et al., 2017). This study revealed that overexpression of NMI suppressed lung cancer cell growth and migration by down-regulating phosphorylated PI3K/AKT without changing the protein levels of PI3K and AKT. Conversely, knockdown of NMI promoted lung cancer cell colony formation and migration by increasing phosphorylated PI3K/AKT without reducing their protein levels. In the case of endometriosis, NMI also suppressed PI3K/AKT signaling in normal endometrium to inhibit their growth upon IFNA exposure. Therefore, the downregulation of NMI activates AKT signaling, enhancing endometriosis progression.

In summary, we propose that endometrial stromal NMI potentially acts as a suppressor of endometriosis. Therefore, the loss of endometrial stromal NMI expression, driven by the ERβ/HDAC8 axis, initiates the progression of endometriosis.

## Methods

### Mice

NOD-SCID (NOD.Cg-Prkdc^scid^/J, Strain number: 001303) female mice (6 weeks old) were obtained from Jackson Laboratory. All mice were housed in the designated animal care facility at Baylor College of Medicine, adhering to the guidelines set by the Institutional Animal Care and Use Committee (IACUC) for the care and use of laboratory animals. All animal experiments in this study were conducted under an IACUC-approved protocol.

### Reagents

Human recombinant IFNα A protein (Sigma-Aldrich catalog number: IF007) was diluted in 0.1% BSA and added to the cells for 24 hours before further analysis.

### Cell culture

We previously isolated and established luciferase-labeled immortalized human endometriotic epithelial cells (IHEEC) and stromal cells (IHESC) from human ovarian endometrioma (Han et al., 2015a, Park, Cho et al., 2022). The cells were cultured in Dulbecco’s Modified Eagle Medium/F-12 (DMEM/F12) supplemented with 10% FBS and maintained in a humidified atmosphere of 5% CO2 and 95% air at 37°C. The medium was replaced every other day.

### Generation of stable NMI knockdown (KD) cell lines

The GIPZ lentiviral shRNA vector targeting human NMI (Clone ID number: V3LHS 323925) was obtained from Open Biosystems. Lentiviral particles were generated in HEK293T cells via transient transfection using the 2nd generation packaging system, which includes the packaging plasmid psPAX2 and envelope plasmid pMD2.G, in conjunction with Lipofectamine 2000 (ThermoFisher, Catalog number: 116680300). The titer of the lentiviral particles was determined using Lenti-X™ GoStix™ Plus (ClonTech, Catalog number: 631280). As a control, a lentivirus containing the non-targeting GIPZ lentiviral shRNA negative control (NT shRNA, Horizon Discover, Catalog number: RHS4348) was used for transduction. Both IHEEC and IHESC were infected with lentiviruses carrying either shNMI or NT shRNA using TransDux MAX™ (System Bioscience, Catalog number: LV860A-1). Subsequently, IHEECs and IHESCs expressing shNMI were selected with 1µg/ml puromycin. The NMI protein levels in NMI KD IHEECs and NMI KD IHESCs were assessed by Western blot analysis using an NMI antibody (Abcam, catalog number: 183724) compared to NT shRNA IHEECs and IHESCs, which served as WT control endometrial cells.

### HDAC8 and NCOR2 knockdown in IHESCs

To knockdown HDAC8, IHESCs were transfected at a final concentration of 10nM with the siHDAC8 duplex or control duplex (Origene, Catalog number: SR311006) using Lipofectamine 2000 (ThermoFisher, Catalog number: 116680300) according to the manufacturer’s instructions. For NCOR2 knockdown, siNCOR2 duplex (Origene, Catalog number: RC212113) was used.

Twenty-four hours post-transfection, the cells were supplemented with fresh DMEM/F12 containing 10% charcoal-stripped FBS. After another 24 hours, total RNA was extracted using RNeasy Plus Mini Kit (Qiagen, catalog number: 74134). The first strand of cDNA was synthesized from 1μg of RNA of each sample using a SuperScript II™ RT kit (Invitrogen, Catalog number: 18064022), according to the manufacturer’s protocol. HDAC8 and NMI expression levels were evaluated using quantitative PCR as described below.

### Growth assay of IHECs NMI KD vs control

Cells were seeded in 96 well plate at the density of 0.03x10^6^ cells/ml. Fresh medium was replaced with either vehicle or 500 or 1000 units/ml IFNα on the following day. After 24 hours of treatment, the cells were treated with premix WST-1 (Takara, Catalog number: MK400) to determine cell viability following manufacturer’s instructions and incubated at 37°C for 1 hour. Their absorbance at 450nm was measured at 620nm by Fisher Scientific accuSkan FC microplate reader. Cell growth was normalized to the vehicle-treated WT.

### Terminal deoxynucleotidyl transferase biotin-dUTP Nick End Labeling (TUNEL) assay

12mm diameter coverslips in 24 well plate were coated with 50µg/ml rat tail Collagen I (Gibco, Catalog number: A10483-01) diluted in 0.02M acetic acid for 1 hour at 37°C. IHESCs and NMI KD IHESCs were seeded on the coverslips. Each group was duplicated. On the next day, 0 or 500 unit/ml IFNA was added, followed by an additional 24-hour incubation. The cells were then subjected to TUNEL assay, using Click-iT™ Plus TUNEL Assay Kits for *In Situ* Apoptosis Detection with Alexa Fluor™ 594 dye (Invitrogen, Catalog number: C10618), as per the manufacturer’s instructions. Nuclei was counterstained with Hoechst 33342 (Sigma, Catalog number: B2883). The stained cells were imaged using Keyence BZ-X800 microscope. Random 6-12 fields of view were captured and counted for TUNEL positive cell percetage.

### Reduction of NMI levels by ERβ in HeLa cells

According to the manufacturer’s instructions, HeLa cells were transfected with ERβ-expressing plasmids (Addgene, Catalog number: 35562) and an empty expression vector (pcDNA 3.1, referred to as MOCK transfection) using Lipofectamine 2000 (ThermoFisher, Catalog number: 116680300). Twenty-four hours post-transfection, the cells were supplemented with phenol red- free DMEM containing 10% charcoal-stripped FBS. After another 24 hours, estradiol (10nM) or a vehicle (ethanol) was added, followed by an additional 24-hour incubation. Total RNA from cells and tissues were isolated using RNeasy Plus Mini Kit (Qiagen, catalog number: 74134).

The first strand of cDNA was synthesized from 1μg of RNA of each sample using a SuperScript II™ RT kit (Invitrogen, Catalog number: 18064022), according to the manufacturer’s protocol. NMI levels were determined by TaqMan probes for NMI (Invitrogen, Catalog number: Hs00190768_m1). Relative mRNA expression was determined using the 2-ΔΔCT method of quantitative PCR, normalized to 18S rRNA levels.

### Development of human endometriotic lesions in NOD-SCID female mice

NOD-SCID female mice, aged six weeks, were implanted with an estrogen pellet (0.36 mg of 17-β estradiol for 60-day release, Innovative Research of America) to stimulate the progression of endometriosis. Following a week of recovery, equal numbers of luciferase-labeled IHEEC (either NMI KD or WT, 1x10^6^ cells) and luciferase-labeled IHESC (either NMI KD or WT, 1x10^6^ cells) were mixed. This mixture was combined with Matrigel® basement membrane matrix (Millipore Sigma, Catalog number: CLS356234) at a 1:1 ratio and then intraperitoneally injected into NOD-SCID mice implanted with an estrogen pellet. After inducing endometriosis in these mice, bioluminescence imaging was determined to visualize human ectopic lesions. This imaging was done using the *In Vivo* Imaging System (IVIS, Perkin Elmer, IVIS® Lumina X5).

### Quantifying bioluminescence data

Mice were anesthetized with a 1.5% isoflurane/air mixture using an Inhalation Anesthesia System (VetEquip). Next, d-Luciferin (ThermoFisher, catalog number: L2916) was intraperitoneally injected at 40 mg/kg mouse body weight. Ten minutes after the D-luciferin injection, the mice were imaged using an IVIS Imaging System (Xenogen) with continuous 1% to 2% isoflurane exposure. Imaging variables were maintained for comparative analysis.

Grayscale-reflected and pseudocolorized images reflecting bioluminescence were superimposed and analyzed using Living Image software (Version 4.4, Xenogen). A region of interest (ROI) was manually selected over the relevant signal intensity regions. The area of the ROI was kept constant across experiments, and the intensity was recorded as total photon counts per second per cm^2^ within the ROI.

### Quantitative PCR (qPCR)

To study relative gene expression, qPCR was conducted using sequence-specific primers in conjunction with the PowerTrack SYBR Green Master Mix (ThermoFisher, catalog number: A46109). Gene-specific primers were designed using the PrimerQuest Tool (https://www.idtdna.com/pages/tools/primerquest). Relative mRNA expression was determined using the 2-ΔΔCT method of quantitative PCR, normalized to GAPDH levels (Livak & Schmittgen, 2001). Each result is represented by at least three technical replicates. Primer sequences were described in Table EV1.

### RNA sequencing analysis

WT control IHESC and NMI KD-IHESCs were cultured in 60mm dishes. Once the cells reached 70-80% confluency, they were incubated with either vehicle or 1000 units/ml IFNα for 24 hours. Each condition was performed in triplicate. Total RNA was extracted from IHESC and NMI KD-IHESCs that were treated with either vehicle or IFNα, using the RNeasy Plus Mini Kit (Qiagen, catalog number: 74134), as per the manufacturer’s instructions. To minimize genomic DNA contamination, the RNase spin column membrane was additionally treated with DNase I (2U/μl). The quality of the total RNA was assessed using the NanoDrop spectrophotometer, Invitrogen Qubit 2.0 quantitation assay, and Agilent Bioanalyzer. Libraries were prepared using the Illumina TruSeq Stranded mRNA library preparation protocol. Sequence reads were trimmed for adapter sequences and low-quality sequences using Galaxy version 23.1.rc1(Community, 2022). Trimmed sequence reads were then mapped to hg38. Subsequently, read count extraction and normalization were conducted on the Galaxy platform.

### Western Blotting

Cells were lysed using a homemade 1x RIPA buffer (150mM NaCl, 50mM Tris pH 8.0, 1% NP- 40, 0.5% Sodium Deoxycholate, 0.1% SDS), supplemented with 1x phosphatase inhibitor cocktails (Gendepot, catalog number: P3200-001) and 1x Xpert protease inhibitor cocktails (Gendepot, catalog number: P3100-001). Samples were incubated on ice for 30 minutes, sonicated for 2 cycles of 10 seconds on and 20 seconds off at 20% power, and then centrifuged for 15 minutes at 15,000 rpm. Equal amounts of proteins were separated by SDS-PAGE and subsequently transferred onto a 0.2 μm PVDF membrane (Cytiva, catalog number: 10-6000-30).

The membranes were blocked with 5% skim milk in TBST for 1 hour at room temperature and then incubated with antigen-specific primary antibodies in 2.5% skim milk in TBST overnight at 4°C. After washing, the membranes were incubated with HRP-tagged secondary antibodies (Abcam, catalog number: ab6721) for 1 hour at room temperature. Signals were visualized using the SuperSignal™ West Pico Plus Chemiluminescent substrate (ThermoFisher, catalog number: 34580). Antibodies used in this experiment were described in Table EV2.

### Formalin-Fixed Paraffin-Embedded (FFPE) for human endometriotic lesions and normal endometrium

Ovarian endometriomas were removed from patients with endometriosis during surgical procedures at Baylor College of Medicine, in accordance with an Institutional Review Board (IRB)-approved human protocol. Normal endometrium was isolated from uteri removed from patients undergoing hysterectomies due to uterine fibroids, also based on an IRB-approved human protocol at Baylor College of Medicine. All patients had refrained from exogenous hormonal treatments for at least three months prior to their surgeries.

Both endometriotic lesions and normal endometrial samples were fixed in 10% buffered formalin phosphate for 24 hours and then stored in 70% EtOH. The tissues were dehydrated with ethanol and xylene using a tissue processor. Following dehydration, the processed tissues were embedded in paraffin.

### Immunohistochemistry of NMI in human endometriotic lesions

Tissues preserved in FFPE were sectioned to a thickness of 7μm. The sliced tissues on the glass slides were deparaffinized in xylene, rehydrated through a gradient of ethanol, and then subjected to immunostaining. Antigen retrieval was carried out using a citrate-based buffer. An antibody against NMI (Abcam, catalog number: 183724) was employed. Specific antigens were visualized using a DAB substrate kit (Vector, catalog number: SK-4100). The intensity of immunostaining was quantified with QuPath software (Bankhead, Loughrey et al., 2017).

### Gene Expression Omnibus (GEO)

*NMI* expression between normal versus endometriotic lesions in human endometriosis patients (GSE25628) were analyzed using Galaxy version 23.1.rc1 (Community, 2022).

### Statistical Analysis

An independent two-tailed Student’s t test was used to assess the statistical significance with GraphPad Prism version 8.0. P< 0.05 was considered statistically significant.

## Abbreviation

Abbreviation: Meaning
ADAM19: ADAM Metallopeptidase Domain 19
AKT: AKT Serine/Threonine Kinase
BSA: Bovine serum albumin
CDH10: Cadherin 10
CDH15: Cadherin 15
ChIP: Chromation ImmunoPrecipitation
COL4A1: Collagen Type IV Alpha 1 Chain
COX-2: Cyclooxygenase 2
Dkk1: Dickkopf 1
DMEM: Dulbecco’s minimum essential medium
E2: Estradiol-17β
ELFN1: Extracellular Leucine Rich Repeat And Fibronectin Type III Domain Containing 1
Erβ: Estrogen Receptor beta
FBS: Fetal bovine serum
FFPE: Formalin-Fixed Paraffin-Embedded
GSK3β: Glycogen Synthase Kinase 3 Beta
HAPLN3: Hyaluronan And Proteoglycan Link Protein 3 IFNa Interferon alpha
IFNAR2: Interferon Alpha And Beta Receptor Subunit 2 IHC Immunohistochemistry
IHEECs: immortalized human endometrial epithelial cells
IHEECs: ERΒ immortalized human endometrial epithelial cells overexpressing ERβ IHESCs immortalized human endometrial stromal cells
IRF9: Interferon Regulatory Factor 9
ISGF3: IFN-stimulated gene factor 3
ISRE: IFN-stimulated responsive element
ITGB3: Integrin Subunit Beta 3
IVIS: In Vivo Imaging System
JAK1: Janus Kinase 1
JUP: Junction Plakoglobin
KD: Knockdown
MFAP5: Microfibril Associated Protein 5
MLKL: mixed lineage kinase domain-like
MMP: Matrix Metallopeptidase
MPZL2: Myelin Protein Zero Like 2
NF-kB: Nuclear Factor Kappa B
NMI: N-Myc and STAT Interactor
NMI KD-IHEECs: NMI knockdown immortalized human endometrial epithelial cells
NMI KD-IHESCs NMI: knockdown immortalized human endometrial stromal cells
NOD-SCID: Nonobese diabetic-severe combined immunodeficiency
NT: Non-Target
PCDH1: Protocadherin 1
PCDHA1: Protocadherin Alpha 1
PGE2: Prostaglandin E2
PI3K: phosphoinositide 3-kinases
PM: promyelocytic leukemia
p-MLKL: phospho-MLKL
p-RIP: phospho-RIP
p-STAT: phospho-STAT
RIP: Receptor-Interacting Serine/Threonine-Protein
SMAD: SMAD Family Member
SORBS1: Sorbin And SH3 Domain Containing 1
SPARC: Secreted Protein Acidic And Cysteine Rich
SPOCK1: SPARC (Osteonectin), Cwcv And Kazal Like Domains Proteoglycan 1
STAT: Signal Transducer And Activator Of Transcription
TBST: Tris-buffered saline
TGF-b: Transforming Growth Factor Beta 1
VCAN: Versican
WT: Wild Type

## Declarations

### Ethics approval and consent to participate

All animal experiments were approved by Institutional Animal Care and Use Committee in Baylor College of Medicine. Experimental protocols and animal care were provided according to the guideline for the care and use of animals established by Baylor College of Medicine.

Ovarian endometriomas were removed from patients with endometriosis during surgical procedures at Baylor College of Medicine following an Institutional Review Board (IRB)- approved human protocol. Normal endometrium was isolated from uteri and removed from patients undergoing hysterectomies due to uterine fibroids, based on an IRB-approved human protocol at Baylor College of Medicine.

### Consent for publication

Not applicable.

### Availability of data and materials

The datasets used and/or analyzed during the current study are available from the corresponding author on reasonable request. RNA sequencing data regarding the INFα response of NMI KD versus control WT immortalized human endometrial stromal cells has been deposited in the GEO database (GSE241751).

### Author contributions

YP conducted major experiments and data analysis. YP and SJH contributed to the design and the writing of the manuscript. XG, provided reagents and critical comments. All authors read and approved the final manuscript.

## Acknowledgements

This project was supported in part by the Genomic and RNA Profiling Core at Baylor College of Medicine with funding from the NIH NCI (P30CA125123) and CPRIT (RP200504) grants.

## References

Acien P, Quereda F, Campos A, Gomez-Torres MJ, Velasco I, Gutierrez M (2002) Use of intraperitoneal interferon alpha-2b therapy after conservative surgery for endometriosis and postoperative medical treatment with depot gonadotropin-releasing hormone analog: a randomized clinical trial. Fertil Steril 78: 705–11

Alam S, Zunic A, Venkat S, Feigin ME, Atanassov BS (2022) Regulation of Cyclin D1 Degradation by Ubiquitin-Specific Protease 27X Is Critical for Cancer Cell Proliferation and Tumor Growth. Molecular cancer research : MCR 20: 1751–1762

Ali S, Mann-Nüttel R, Schulze A, Richter L, Alferink J, Scheu S (2019) Sources of Type I Interferons in Infectious Immunity: Plasmacytoid Dendritic Cells Not Always in the Driver’s Seat. Frontiers in immunology 10: 778

Balasubramanian S, Ramos J, Luo W, Sirisawad M, Verner E, Buggy JJ (2008) A novel histone deacetylase 8 (HDAC8)-specific inhibitor PCI-34051 induces apoptosis in T-cell lymphomas. Leukemia 22: 1026–34

Bankhead P, Loughrey MB, Fernández JA, Dombrowski Y, McArt DG, Dunne PD, McQuaid S, Gray RT, Murray LJ, Coleman HG, James JA, Salto-Tellez M, Hamilton PW (2017) QuPath: Open source software for digital pathology image analysis. Scientific reports 7: 16878

Bulun SE (2009) Endometriosis. N Engl J Med 360: 268–79

Bulun SE, Cheng YH, Pavone ME, Xue Q, Attar E, Trukhacheva E, Tokunaga H, Utsunomiya H, Yin P, Luo X, Lin Z, Imir G, Thung S, Su EJ, Kim JJ (2010) Estrogen receptor-beta, estrogen receptor-alpha, and progesterone resistance in endometriosis. Semin Reprod Med 28: 36–43

Chawla-Sarkar M, Lindner DJ, Liu YF, Williams BR, Sen GC, Silverman RH, Borden EC (2003) Apoptosis and interferons: Role of interferon-stimulated genes as mediators of apoptosis. Apoptosis 8: 237-249

Chen HP, Zhao YT, Zhao TC (2015) Histone deacetylases and mechanisms of regulation of gene expression. Critical reviews in oncogenesis 20: 35–47

Community TG (2022) The Galaxy platform for accessible, reproducible and collaborative biomedical analyses: 2022 update. Nucleic Acids Research 50: W345–W351

Dicitore A, Castiglioni S, Saronni D, Gentilini D, Borghi MO, Stabile S, Vignali M, Di Blasio AM, Persani L, Vitale G (2018) Effects of human recombinant type I IFNs (IFN-alpha2b and IFN-beta1a) on growth and migration of primary endometrial stromal cells from women with deeply infiltrating endometriosis: A preliminary study. Eur J Obstet Gynecol Reprod Biol 230: 192–198

Duong V, Licznar A, Margueron R, Boulle N, Busson M, Lacroix M, Katzenellenbogen BS, Cavaillès V, Lazennec G (2006) ERα and ERβ expression and transcriptional activity are differentially regulated by HDAC inhibitors. Oncogene 25: 1799–1806

Fillmore RA, Mitra A, Xi Y, Ju J, Scammell J, Shevde LA, Samant RS (2009) Nmi (N-Myc interactor) inhibits Wnt/beta-catenin signaling and retards tumor growth. International journal of cancer 125: 556–64

Halme J, Hammond MG, Hulka JF, Raj SG, Talbert LM (1984) Retrograde menstruation in healthy women and in patients with endometriosis. Obstet Gynecol 64: 151–4

Han SJ, Jung SY, Wu SP, Hawkins SM, Park MJ, Kyo S, Qin J, Lydon JP, Tsai SY, Tsai MJ, DeMayo FJ, O’Malley BW (2015a) Estrogen Receptor beta Modulates Apoptosis Complexes and the Inflammasome to Drive the Pathogenesis of Endometriosis. Cell 163: 960–74

Han SJ, Jung SY, Wu SP, Hawkins SM, Park MJ, Kyo S, Qin J, Lydon JP, Tsai SY, Tsai MJ, DeMayo FJ, O’Malley BW (2015b) Estrogen Receptor β Modulates Apoptosis Complexes and the Inflammasome to Drive the Pathogenesis of Endometriosis. Cell 163: 960–74

Han SJ, Lee JE, Cho YJ, Park MJ, O’Malley BW (2019a) Genomic Function of Estrogen Receptor beta in Endometriosis. Endocrinology 160: 2495–2516

Han SJ, Lee JE, Cho YJ, Park MJ, O’Malley BW (2019b) Genomic Function of Estrogen Receptor β in Endometriosis. Endocrinology 160: 2495–2516

He T, Qiao Y, Yang Q, Chen J, Chen Y, Chen X, Hao Z, Lin M, Shao Z, Wu P, Xu F (2022) NMI: a potential biomarker for tumor prognosis and immunotherapy. Frontiers in pharmacology 13: 1047463

Herzer K, Hofmann TG, Teufel A, Schimanski CC, Moehler M, Kanzler S, Schulze-Bergkamen H, Galle PR (2009) IFN-alpha-induced apoptosis in hepatocellular carcinoma involves promyelocytic leukemia protein and TRAIL independently of p53. Cancer Res 69: 855–62

Ingelmo JM, Quereda F, Acien P (1999) Intraperitoneal and subcutaneous treatment of experimental endometriosis with recombinant human interferon-alpha-2b in a murine model. Fertil Steril 71: 907–11

Kao LC, Germeyer A, Tulac S, Lobo S, Yang JP, Taylor RN, Osteen K, Lessey BA, Giudice LC (2003) Expression profiling of endometrium from women with endometriosis reveals candidate genes for disease-based implantation failure and infertility. Endocrinology 144: 2870–81

Klinge CM, Jernigan SC, Mattingly KA, Risinger KE, Zhang J (2004) Estrogen response element-dependent regulation of transcriptional activation of estrogen receptors alpha and beta by coactivators and corepressors. Journal of molecular endocrinology 33: 387–410

Lebrun SJ, Shpall RL, Naumovski L (1998) Interferon-induced upregulation and cytoplasmic localization of Myc-interacting protein Nmi. J Interferon Cytokine Res 18: 767–71

Li L, Chen SN, Wang KL, Li N, Pang AN, Liu LH, Li B, Hou J, Wang S, Nie P (2023) Interaction of Nmi and IFP35 Promotes Mutual Protein Stabilization and IRF3 and IRF7 Degradation to Suppress Type I IFN Production in Teleost Fish. Journal of immunology (Baltimore, Md : 1950) 210: 1494-1507

Li WX (2008) Canonical and non-canonical JAK-STAT signaling. Trends Cell Biol 18: 545–51

Livak KJ, Schmittgen TD (2001) Analysis of relative gene expression data using real-time quantitative PCR and the 2(-Delta Delta C(T)) Method. Methods (San Diego, Calif) 25: 402-8

Lu Y, Nie J, Liu X, Zheng Y, Guo SW (2010) Trichostatin A, a histone deacetylase inhibitor, reduces lesion growth and hyperalgesia in experimentally induced endometriosis in mice. Human reproduction (Oxford, England) 25: 1014-25

Lyons SD, Chew SS, Thomson AJ, Lenart M, Camaris C, Vancaillie TG, Abbott JA (2006) Clinical and quality-of-life outcomes after fertility-sparing laparoscopic surgery with bowel resection for severe endometriosis. J Minim Invasive Gynecol 13: 436–41

Madanes D, Bilotas MA, Bastón JI, Singla JJ, Meresman GF, Barañao RI, Ricci AG (2020) PI3K/AKT pathway is altered in the endometriosis patient’s endometrium and presents differences according to severity stage. Gynecological endocrinology : the official journal of the International Society of Gynecological Endocrinology 36: 436–440

Majali-Martinez A, Hoch D, Tam-Amersdorfer C, Pollheimer J, Glasner A, Ghaffari-Tabrizi- Wizsy N, Beristain AG, Hiden U, Dieber-Rotheneder M, Desoye G (2020) Matrix metalloproteinase 15 plays a pivotal role in human first trimester cytotrophoblast invasion and is not altered by maternal obesity. FASEB journal : official publication of the Federation of American Societies for Experimental Biology 34: 10720–10730

Marineau A, Khan KA, Servant MJ (2020) Roles of GSK-3 and beta-Catenin in Antiviral Innate Immune Sensing of Nucleic Acids. Cells 9

Matsuzaki S, Canis M, Pouly JL, Botchorishvili R, Déchelotte PJ, Mage G (2006) Differential expression of genes in eutopic and ectopic endometrium from patients with ovarian endometriosis. Fertility and sterility 86: 548–53

Mazewski C, Perez RE, Fish EN, Platanias LC (2020) Type I Interferon (IFN)-Regulated Activation of Canonical and Non-Canonical Signaling Pathways. Front Immunol 11: 606456

McComb S, Cessford E, Alturki NA, Joseph J, Shutinoski B, Startek JB, Gamero AM, Mossman KL, Sad S (2014) Type-I interferon signaling through ISGF3 complex is required for sustained Rip3 activation and necroptosis in macrophages. Proc Natl Acad Sci U S A 111: E3206–13

Meng D, Chen Y, Yun D, Zhao Y, Wang J, Xu T, Li X, Wang Y, Yuan L, Sun R, Song X, Huai C, Hu L, Yang S, Min T, Chen J, Chen H, Lu D (2015) High expression of N-myc (and STAT) interactor predicts poor prognosis and promotes tumor growth in human glioblastoma. Oncotarget 6: 4901–19

Mohankumar K, Li X, Sung N, Cho YJ, Han SJ, Safe S (2020) Bis-Indole-Derived Nuclear Receptor 4A1 (NR4A1, Nur77) Ligands as Inhibitors of Endometriosis. Endocrinology 161

Monnin N, Fattet AJ, Koscinski I (2023) Endometriosis: Update of Pathophysiology, (Epi) Genetic and Environmental Involvement. Biomedicines 11

Monsivais D, Dyson MT, Yin P, Coon JS, Navarro A, Feng G, Malpani SS, Ono M, Ercan CM, Wei JJ, Pavone ME, Su E, Bulun SE (2014) ERbeta- and prostaglandin E2-regulated pathways integrate cell proliferation via Ras-like and estrogen-regulated growth inhibitor in endometriosis. Mol Endocrinol 28: 1304–15

Newby BN, Brusko TM, Zou B, Atkinson MA, Clare-Salzler M, Mathews CE (2017) Type 1 Interferons Potentiate Human CD8(+) T-Cell Cytotoxicity Through a STAT4- and Granzyme B- Dependent Pathway. Diabetes 66: 3061–3071

Park Y, Cho YJ, Sung N, Park MJ, Guan X, Gibbons WE, O’Malley BW, Han SJ (2022) Oleuropein suppresses endometriosis progression and improves the fertility of mice with endometriosis. J Biomed Sci 29: 100

Park Y, Han SJ (2022) Interferon Signaling in the Endometrium and in Endometriosis. Biomolecules 12

Pazhohan A, Amidi F, Akbari-Asbagh F, Seyedrezazadeh E, Farzadi L, Khodarahmin M, Mehdinejadiani S, Sobhani A (2018) The Wnt/β-catenin signaling in endometriosis, the expression of total and active forms of β-catenin, total and inactive forms of glycogen synthase kinase-3β, WNT7a and DICKKOPF-1. Eur J Obstet Gynecol Reprod Biol 220: 1-5

Pia MM, Anna LV, Ulla BK (2012) Virus Infection and Type I Interferon in Endometriosis. In Endometriosis, Koel C, Baidyanath C (eds) p Ch. 12. Rijeka: IntechOpen

Pruitt HC, Devine DJ, Samant RS (2016) Roles of N-Myc and STAT interactor in cancer: From initiation to dissemination. Int J Cancer 139: 491–500

Sampson JA (1925) Heterotopic or misplaced endometrial tissue. American Journal of Obstetrics and Gynecology 10: 649–664

Santulli P, Marcellin L, Tosti C, Chouzenoux S, Cerles O, Borghese B, Batteux F, Chapron C (2015) MAP kinases and the inflammatory signaling cascade as targets for the treatment of endometriosis? Expert opinion on therapeutic targets 19: 1465–83

Sarhan J, Liu BC, Muendlein HI, Weindel CG, Smirnova I, Tang AY, Ilyukha V, Sorokin M, Buzdin A, Fitzgerald KA, Poltorak A (2019) Constitutive interferon signaling maintains critical threshold of MLKL expression to license necroptosis. Cell Death & Differentiation 26: 332–347

Saunders PTK, Horne AW (2021) Endometriosis: Etiology, pathobiology, and therapeutic prospects. Cell 184: 2807–2824

Shi W, Yao X, Fu Y, Wang Y (2022) Interferon-alpha and its effects on cancer cell apoptosis. Oncol Lett 24: 235

Shi WY, Cao C, Liu L (2016) Interferon alpha Induces the Apoptosis of Cervical Cancer HeLa Cells by Activating both the Intrinsic Mitochondrial Pathway and Endoplasmic Reticulum Stress-Induced Pathway. Int J Mol Sci 17

Sourial S, Tempest N, Hapangama DK (2014) Theories on the pathogenesis of endometriosis. International journal of reproductive medicine 2014: 179515

Suen JL, Chang Y, Shiu YS, Hsu CY, Sharma P, Chiu CC, Chen YJ, Hour TC, Tsai EM (2019) IL-10 from plasmacytoid dendritic cells promotes angiogenesis in the early stage of endometriosis. The Journal of pathology 249: 485–497

Sugihara K, Kobayashi Y, Suzuki A, Tamura N, Motamedchaboki K, Huang CT, Akama TO, Pecotte J, Frost P, Bauer C, Jimenez JB, Nakayama J, Aoki D, Fukuda MN (2014) Development of pro-apoptotic peptides as potential therapy for peritoneal endometriosis. Nature communications 5: 4478

Tosti C, Biscione A, Morgante G, Bifulco G, Luisi S, Petraglia F (2017) Hormonal therapy for endometriosis: from molecular research to bedside. Eur J Obstet Gynecol Reprod Biol 209: 61–66

Vanden Berghe T, Linkermann A, Jouan-Lanhouet S, Walczak H, Vandenabeele P (2014) Regulated necrosis: the expanding network of non-apoptotic cell death pathways. Nat Rev Mol Cell Biol 15: 135–47

Wang J, Zou K, Feng X, Chen M, Li C, Tang R, Xuan Y, Luo M, Chen W, Qiu H, Qin G, Li Y, Zhang C, Xiao B, Kang L, Kang T, Huang W, Yu X, Wu X, Deng W (2017) Downregulation of NMI promotes tumor growth and predicts poor prognosis in human lung adenocarcinomas. Molecular cancer 16: 158

Wu G, Huang H, Garcia Abreu J, He X (2009) Inhibition of GSK3 phosphorylation of beta- catenin via phosphorylated PPPSPXS motifs of Wnt coreceptor LRP6. PloS one 4: e4926

Wu Y, Starzinski-Powitz A, Guo SW (2007) Trichostatin A, a histone deacetylase inhibitor, attenuates invasiveness and reactivates E-cadherin expression in immortalized endometriotic cells. Reproductive sciences (Thousand Oaks, Calif) 14: 374-82

Xu X, Chai K, Chen Y, Lin Y, Zhang S, Li X, Qiao W, Tan J (2018) Interferon activates promoter of Nmi gene via interferon regulator factor-1. Mol Cell Biochem 441: 165–171

Xue Q, Lin Z, Cheng YH, Huang CC, Marsh E, Yin P, Milad MP, Confino E, Reierstad S, Innes J, Bulun SE (2007) Promoter methylation regulates estrogen receptor 2 in human endometrium and endometriosis. Biol Reprod 77: 681–7

Yang W, Feng Y, Zhou J, Cheung OK, Cao J, Wang J, Tang W, Tu Y, Xu L, Wu F, Tan Z, Sun H, Tian Y, Wong J, Lai PB, Chan SL, Chan AW, Tan PB, Chen Z, Sung JJ et al. (2021) A selective HDAC8 inhibitor potentiates antitumor immunity and efficacy of immune checkpoint blockade in hepatocellular carcinoma. Science translational medicine 13

Zakhari A, Delpero E, McKeown S, Tomlinson G, Bougie O, Murji A (2021) Endometriosis recurrence following post-operative hormonal suppression: a systematic review and meta- analysis. Hum Reprod Update 27: 96–107

Zheng H, Liu X, Guo SW (2023a) Aberrant expression of histone deacetylase 8 in endometriosis and its potential as a therapeutic target. Reprod Med Biol 22: e12531

Zheng H, Liu X, Guo SW (2023b) Corroborating evidence for aberrant expression of histone deacetylase 8 in endometriosis. Reproductive medicine and biology 22: e12527

Zhu M, John S, Berg M, Leonard WJ (1999) Functional association of Nmi with Stat5 and Stat1 in IL-2- and IFNgamma-mediated signaling. Cell 96: 121–30

Zondervan KT, Becker CM, Missmer SA (2020) Endometriosis. N Engl J Med 382: 1244–1256

